# No evidence for disruption of global patterns of nest predation in shorebirds

**DOI:** 10.1101/601047

**Authors:** Martin Bulla, Jeroen Reneerkens, Emily L. Weiser, Aleksandr Sokolov, Audrey R. Taylor, Benoît Sittler, Brian J. McCaffery, Dan R. Ruthrauff, Daniel H. Catlin, David C. Payer, David H. Ward, Diana V. Solovyeva, Eduardo S.A. Santos, Eldar Rakhimberdiev, Erica Nol, Eunbi Kwon, Glen S. Brown, Glenda D. Hevia, H. River Gates, James A. Johnson, Jan A. van Gils, Jannik Hansen, Jean-François Lamarre, Jennie Rausch, Jesse R. Conklin, Joe Liebezeit, Joёl Bêty, Johannes Lang, José A. Alves, Juan Fernández-Elipe, Klaus Michael-Exo, Loïc Bollache, Marcelo Bertellotti, Marie-Andrée Giroux, Martijn van de Pol, Matthew Johnson, Megan L. Boldenow, Mihai Valcu, Mikhail Soloviev, Natalya Sokolova, Nathan R. Senner, Nicolas Lecomte, Nicolas Meyer, Niels Martin Schmidt, Olivier Gilg, Paul A. Smith, Paula Machín, Rebecca L. McGuire, Ricardo A.S. Cerboncini, Richard Ottvall, Rob S.A. van Bemmelen, Rose J. Swift, Sarah T. Saalfeld, Sarah E. Jamieson, Stephen Brown, Theunis Piersma, Tomas Albrecht, Verónica D’Amico, Richard B. Lanctot, Bart Kempenaers

**Affiliations:** Max Planck Institute for Ornithology, Department of Behavioural Ecology & Evolutionary Genetics, E Gwinner Str., 82319 Seewiesen, Germany; NIOZ Royal Netherlands Institute for Sea Research, Department of Coastal Systems and Utrecht University, PO Box 59, 1790 AB Den Burg, Texel, The Netherlands; Czech University of Life Sciences Prague, Faculty of Environmental Sciences, Kamýcká 129, 16521 Prague, Czech Republic; Groningen Institute for Evolutionary Life Sciences (GELIFES), University of Groningen, Conservation Ecology Group, P.O. Box 11103, 9700 CC Groningen, The Netherlands; U.S. Geological Survey, Upper Midwest Environmental Sciences Center, 2630 Fanta Reed Rd, La Crosse, WI 54603, U.S.A.; Institute of Plant and Animal Ecology, Arctic research station, Zelenaya Gorka, 21, 629400, Labytnangi, Russia; University of Alaska Anchorage, Department of Geography and Environmental Studies, 3211 Providence Drive, 99508, Anchorage, U.S.A; University of Freiburg, Nature Conservation and Landscape Ecology, Tennenbacher Str. 4, 79106, Freiburg, Germany; Arctic Ecology Research Group (GREA), 16 rue de Vernot, F-21440, Francheville, France; US Fish and Wildlife Service, Yukon Delta National Wildlife Refuge, 53980 County Highway D, 54839, Grand View, U.S.A; US Geological Survey, Alaska Science Center, 4210 University Dr., 99508, Anchorage, U.S.A; Virginia Tech, Department of Fish and Wildlife Conservation, 310 West Campus Drive, 24061, Blacksburg, U.S.A; National Park Service, Natural Resource Sciences, 240 W 5th Ave., 99501, Anchorage, U.S.A; Institute of Biological Problems of the North, FEB RAS, 18 Portovaya, Magadan, 685000, Russia; Universidade de São Paulo, BECO do Departamento de Zoologia, Rua do Matão, Trav. 14, n° 101, 05508-090, São Paulo, Brazil; Lomonosov Moscow State University, Department of Vertebrate Zoology, Leninskiye Gory 1/12, 119234, Moscow, Russia; Trent University, Biology Department, 2140 East Bank Drive, K9J 7B8, Peterborough, Canada; Ministry of Natural Resources and Forestry, Wildlife Research and Monitoring, 2140 East Bank Drive, K9L 1Z8, Peterborough, Canada; Centro para el Estudio de Sistemas Marinos (CESIMAR)-CCT CONICET-CENPAT, Grupo de Ecofisiología Aplicada al Manejo y Conservación de Fauna Silvestre, Bvd. Brown 2915, 9120, Puerto Madryn, Argentina; National Audubon Society, Pacific Flyway Program, 431 W 7th Ave #101, 99501, Anchorage, U.S.A; U.S. Fish and Wildlife Service, Migratory Bird Management, 1011 East Tudor Road, 99503, Anchorage, U.S.A; Aarhus University, Department of Bioscience, Frederiksborgvej 399, 4000, Roskilde, Denmark; Polar Knowledge Canada, Science & Technology Program, 1 Uvajuq Place, X0B0C0, Cambridge Bay, Canada; Canadian Wildlife Service, Environment and Climate Change Canada, P.O. Box 2310, X1A 2P7, Yellowknife, Canada; Audubon Society of Portland, 5151 NW Cornell Road, 97210, Portland, U.S.A; University of Quebec at Rimouski, Department of Biology and Center for Northern Studies, 300 Allee des ursulines, G5L 3A1, Rimouski, Canada; Giessen University, Clinic for Birds, Reptiles, Amphibians and Fish / Working Group for Wildlife Biology, Frankfurter Str. 91-93, 35392, Giessen, Germany; DBIO & CESAM-Centre for Environmental and Marine Studies, Department of Biology, University of Aveiro, 3810-193, Aveiro, Portugal; South Iceland Research Centre, University of Iceland, Fjolheimar IS-800 Selfoss & IS-861 Gunnarsholt, Iceland; Aptdo.correos 32, 5480, Candeleda, Spain; Institute of Avian Research “Vogelwarte Helgoland”, An der Vogelwarte 21, 26386, Wilhelmshaven, Germany; Université de Franche-Comté, Laboratoire Chrono-environnement, UMR 6249 CNRS-UFC, F-25000, Besançon, France; Université de Moncton, Faculty of Sciences, K.-C.-Irving Research Chair in Environmental Sciences and Sustainable Development, 18 avenue Antonine-Maillet, E1A 3E9, Moncton, Canada; Netherlands Institute of Ecology (NIOO-KNAW), Department of Animal Ecology, Droevendaalsesteeg 10, 6708PB, Wageningen, the Netherlands; USDA Forest Service, Plumas National Forest, 159 Lawrence Street, 95971, Quincy, U.S.A; University of Alaska Fairbanks, Biology and Wildlife Department, 2090 Koyukuk Drive, 99775, Fairbanks, U.S.A; University of South Carolina, Department of Biological Sciences, 715 Sumter Street, 29208, Columbia, U.S.A; Uninversité de Moncton, Department of Biology and Canada Research Chair in Polar and Boreal Ecology, 18, Antonine-Maillet, E1A 3E9, Moncton, Canada; Aarhus University, Arctic Research Centre, Ny Munkegade 114, 8000, Aarhus C, Denmark; Environment and Climate Change Canada, National Wildlife Research Centre, 1125 Colonel By Dr, K1S 5B6, Ottawa, Canada; Wildlife Conservation Society, Arctic Beringia Program, 3550 Airport Way, Unit 5, 99709, Fairbanks, U.S.A; Universidade Federal do Paraná, Departamento de Zoologia, Av. Coronel Francisco H. dos Santos 100, 81531-980, Curitiba, Brazil; Frostavallsv 325, S-24393, Hoor, Sweden; Wageningen Marine Research, Haringkade 1, 1976CP, IJmuiden, the Netherlands; Cornell University, Cornell Lab of Ornithology and Department of Natural Resources, 159 Sapsucker Woods Rd, 14850, Ithaca, U.S.A; Simon Fraser University, Centre for Wildlife Ecology, 8888 University Dr, V5A1S6, Burnaby, Canada; Manomet Inc., Shorebird Recovery Program, PO Box 545, 05154, Saxtons River, U.S.A; Czech Academy of Sciences, Institute of Vertebrate Biology, Kvetna 8, 60300, Brno, Czech Republic; Charles University, Faculty of Science, 128 44 Prague, Czech Republic

## Abstract

Kubelka et al. (Science, 9 November 2018, p. 680-683) claim that climate change has disrupted patterns of nest predation in shorebirds. They report that predation rates have increased since the 1950s, especially in the Arctic. We describe methodological problems with their analyses and argue that there is no solid statistical support for their claims.

---

Climate change affects organisms in a variety of ways (*1–4*), including through changes in interactions between species. A recent study (*5*, referred to as “the Authors”) reports that a specific type of trophic interaction, namely depredation of shorebird nests, increased globally over the past 70 years. The Authors state that their results are “consistent with climate - induced shifts in predator-prey relationships”. They also claim that the historical perception of a latitudinal gradient in nest predation, with the highest rates in the tropics, “has been recently reversed in the Northern Hemisphere, most notably in the Arctic.” They conclude that “the Arctic now represents an extensive ecological trap… for migrating birds, with a predicted negative impact on their global population dynamics”. These conclusions have far-reaching implications, for evolutionary and population ecology, and for shorebird conservation and related policy decisions (*6*). Therefore, such claims require robust evidence, strongly supported by the data. Here we dispute this evidence.

First, the Authors graphically show non-linear, spatio-temporal variation in predation rates (their Fig. 2AB and 3), and suggest that in recent years, predation has strongly increased in North temperate and especially Arctic regions, but less so in other areas. However, they only statistically test for linear changes in predation rates over time for all regions combined, and for each geographical region (their Table S2) or period (before- and after-2000; their Table S6) separately. To substantiate their conclusions, the Authors should have presented statistical evidence for an interaction between region/latitude and year/period on predation rate. Moreover, their analyses control for spatial auto-correlation, but failed to model non-independence of data from the same site (pseudo-replication).

Using the Authors’ data, we ran a set of mixed-effect models, structurally reflecting their results depicted in their Fig. 2AB and 3, but including location as a random factor (Table 1, (*7*)). These analyses show (a) that much of the variation in nest predation rate is explained by study site (>60%, compared to species: <5%), implying a reduced effective sample size, (b) that all regions – except the South temperate – show similar predation rates, and (c) that nest predation rates increase over time similarly across all geographical areas (Fig. 1A-F). Linear models without interaction terms are much better supported than non-linear models with interactions (Table 1), indicating that predation rates in the Arctic are not increasing any faster than elsewhere (Fig. 1BCEF). Thus, these results provide no evidence that the rate at which nest predation increased over time varies geographically.

**Figure 1.**
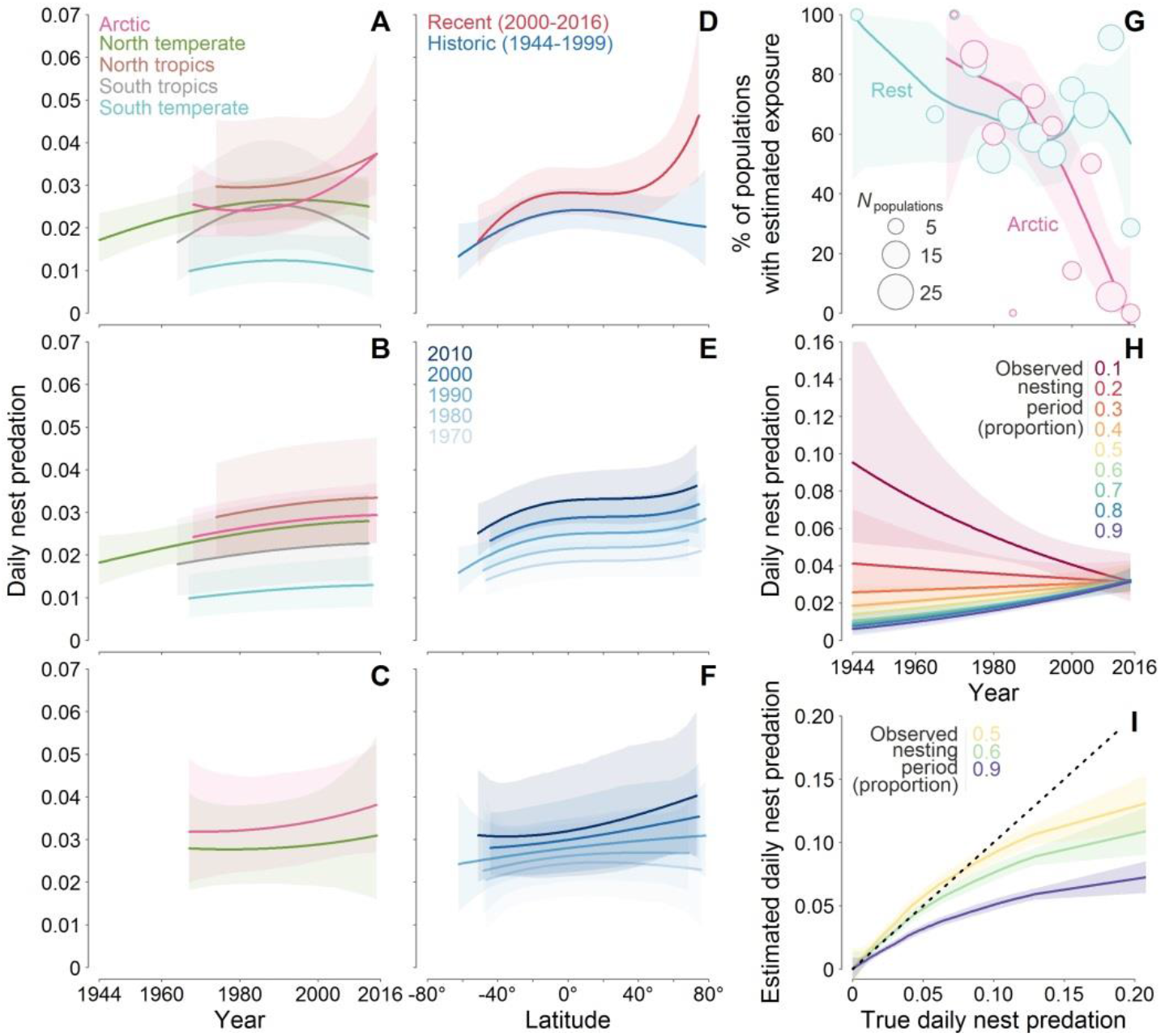
Spatio-temporal variation in daily nest predation rates of shorebirds. (**A-C**) Predation rate in relation to year for different geographical regions; with interaction and using all populations (**A**), without interaction and using all populations (**B**), with interaction and using only the 88 populations with known exposure from the Arctic and North temperate region (**C**). The model behind (**A**) is ~18 times less supported by the data than the model behind (**B**) (Table 1). (**D-F**) Predation rate in relation to latitude for different periods: with interaction (period as two-level factor) and using all populations (**D**), without interaction (year as continuous variable) and using all populations (**E**), with interaction and using only the 98 populations with known exposure (**F**). The model behind (**D**) is ~70 times less supported than the model behind (**E**) (Table 1). (**A-F, H**) Lines and shaded areas represent model predictions with 95% confidence intervals based on posterior distribution of 5,000 simulated values. Note the weak (P>0.64) temporal increase in (**C**) (estimate [95%CI] = 0.08 [-0.07 – 0.2] from a linear model without interaction) and (**F**) (0.06 [-0.09 – 0.17]). See Table 1 for model description and comparison and (7) for details. (**G**) Temporal change in the percentage of populations in which exposure was estimated (following (*10*)) to calculate predation rate. Note the sharp decline in the Arctic compared to the other regions (for overall and region-specific changes, see (*7*)). (**H**) Modeled changes in predation rate over time assuming different values of **B** (proportion of nesting period observed; higher values indicate nests found sooner after egg laying) for populations with unknown exposure and year <2000 (leaving the original estimates for all remaining populations). This exercise explores the sensitivity of the results to using older studies where the stage at which nests were found is less certain. (**I**) Relationship between true and estimated predation rate for different values of *B* (*N* = 65 populations, as in (*5*)). The dashed line indicates a slope of one, i.e. estimated values equaling true values. (**G, I**) Lines and shaded areas represent locally estimated scatterplot smoothing with 95% confidence intervals. Circles in (**G**) represent data for 5-year intervals.

**Table 1.**
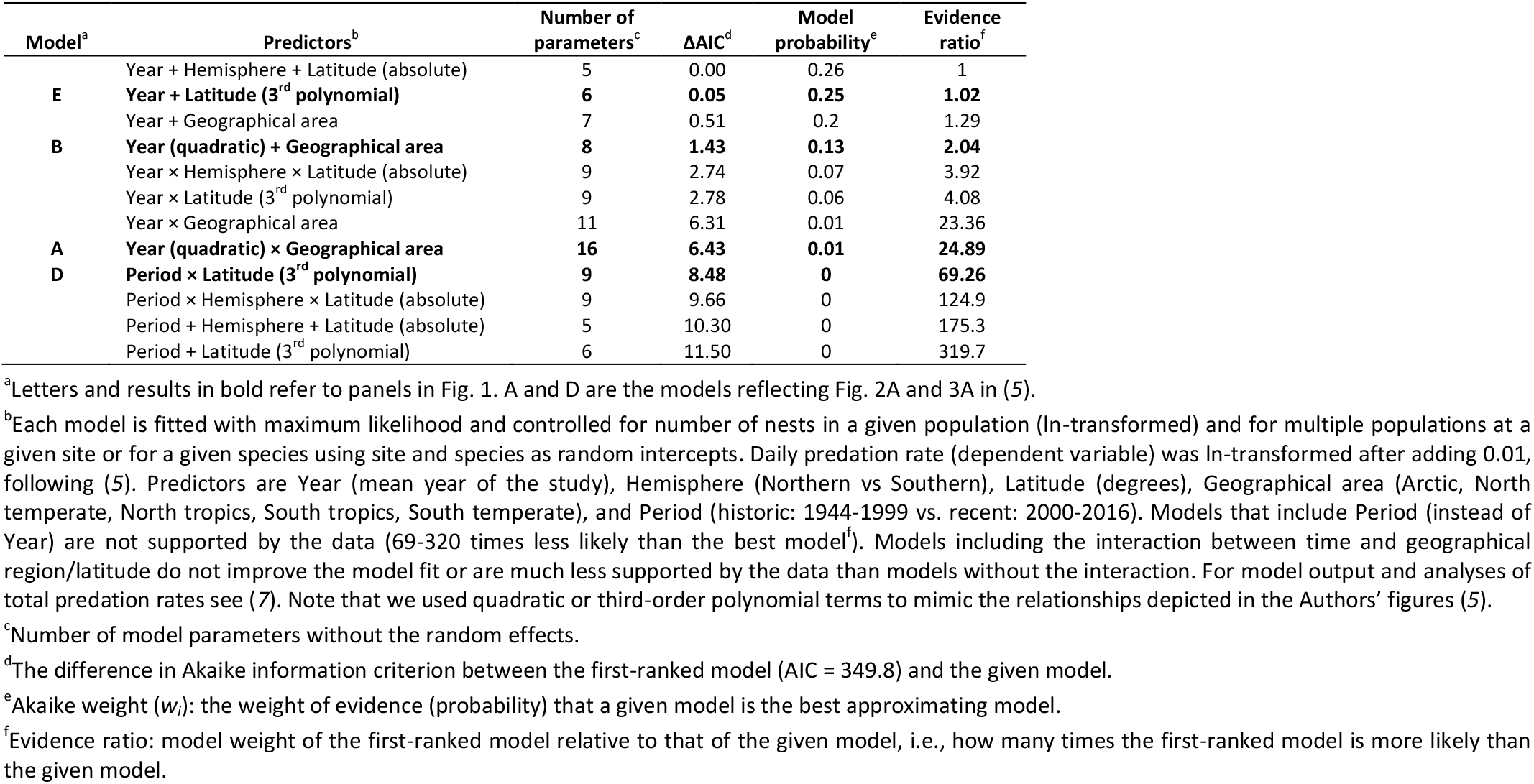
Comparison of models explaining spatio-temporal variation in daily nest predation rate using the Authors’ original data.

Second, for the period under study, not only the climate has changed, but also the research methods. Hence, it remains unclear whether nest predation rates have indeed increased over time and if so, why. The Authors used the Mayfield method (*8, 9*) to calculate daily nest predation rates, as the number of depredated nests divided by “exposure” (the total time all nests were observed in days). However, 59% of the 237 populations used by the Authors lacked information on exposure. The Authors circumvented this problem by estimating exposure based on the description of nest search intensity in the respective studies (10). The key question is when nests were found. The Authors decided that in 114 populations, nests were found such that 60% of the nesting period (egg laying and incubation combined) was “observed” (*B*=0.6; nests searched once or twice a week). For 14 populations they used *B*=0.9 (nests searched daily or found just after laying) and for 11 populations *B*=0.5 (assuming nest found mid-way during the nesting period). However, the choice of *B*-value remains subjective (*7*) and for 38% of the 128 populations where the Authors used *B*>0.5, we found no information in the reference to suggest this was appropriate. This issue is not trivial, because using higher *B*-values, i.e., assuming that nests were found earlier than they actually were, *overestimates* exposure and hence *underestimates* nest predation rates. Importantly, the proportion of populations with estimated exposure declines over time (*7*), particularly after 2000 and especially in the Arctic (Fig. 1G). The timing of the decline coincides with the Authors’ definition of historic and recent data and with the suggested exponential rise of predation in the Arctic (their Fig. 2AB and 3AB). Indeed, the results are sensitive to variation in estimated exposure during the “historic period” (Fig. 1H). Although the Authors correctly state that the estimated and true predation rates are highly correlated (using studies with quantitative information on exposure; see supplementary material in (5)), the true rate is typically underestimated for the higher *B*-values used by the Authors (Fig. 1I). Given these issues, the main result – i.e. the apparent increase in daily nest predation rate over time, especially in the Arctic – may simply be an artifact. To further assess the robustness of the change in predation rate over time, we used only populations where nest predation rates were calculated based on known exposure (*N*=98). These analyses reduced the effect of year by ~50% (*7*) and resulted in weak, non-significant linear trends (Fig. 1CF), suggesting that there is little evidence for changing predation rates.

Finally, we note that nest searching effort and frequency of nest visits likely increased in recent years as researchers learned how best to obtain accurate estimates of nest survival (*11–13*). Researchers also intensified their activities, e.g. capturing adults to band, tag and collect samples, and placing monitoring equipment near nests, which may increase predation rate (*14–15*). Thus, an increase in the quality of data reporting as well as increased research activity around nests may have further induced a time-dependent bias in estimates with an underestimation of true predation rates in the historic data (see above), and perhaps an overestimation in the contemporary data.

In summary, re-analysis of the Authors’ data, evaluation of the quality and interpretation of the published data used, and considerations about changes in research methods over the past 70 years lead us to conclude that there is no robust evidence for a global disruption of nest predation rates due to climate change. We argue that the Authors’ claim that the Arctic has become an ecological trap for breeding shorebirds is untenable.

## Acknowledgments

We thank all that contributed to the original studies on predation, shared data, helped finding the original data or helped in preparing this Technical Comment (for details see (*7*)).

## Author contributions

MB, JR, RL and BK initiated this work. MB, JR, EW, DS, RL, and BK extracted and evaluated the data; most authors collected data contained in the primary sources. EW investigated the differences between newly extracted and original data. MB conducted the statistical analyses and drew the figure with help from MV and EW and input from JR, RL and BK. BK and MB wrote the manuscript with input from JR, EW and RL, and finalized it with input from all authors.

## Data and materials availability

All data, code, methods, results, figures and tables associated with this commentary are freely available from Open Science Framework (*7*).

## SUPPORTING INFORMATION

### General statistical procedures

R version 3.5.1^1^ was used for all statistical analyses, the ‘coxme’ R package^2^ for replicating Kubelka et al.’s^3^ models and the ‘Ime4’ R package^4^ for fitting all other mixed-effect models. We used the ‘sim’ function from the ‘arm’ R package and non-informative prior-distribution^5, 6^ to create a sample of 5,000 simulated values for each model parameter (i.e. posterior distribution). We report effect sizes and model predictions by the medians, and the uncertainty of the estimates and predictions by the Bayesian 95% credible intervals represented by 2.5 and 97.5 percentiles (95%CI) from the posterior distribution of the 5,000 simulated or predicted values. We estimated the variance components with the ‘lmer’ function from the ‘lme4’ R package^4^. The models were fitted with restricted maximum likelihood and controlled for number of nests (ln-transformed). Following Kubelka et al.’s procedure, dependent variable ‘daily predation rate’ was ln-transformed (after adding 0.01) and ‘total predation rate’ was left as a proportion. We have checked whether the assumptions of all models were met (see the online material).

In all model comparisons we assessed the model fit by Akaike’s Information Criterion using maximum likelihood and the ‘AlC’ function in R^7^.

### Testing global patterns

#### Geographical zones

Using Kubelka et al.’s data and model (see their Table S2A), we first tested for the difference in patterns of predation rates between the geographical zones by testing for the interaction between ‘mean year’ of the study and five ‘geographical zones’ (Table S1A). We also specified a similar model, but with widely used ‘lmer’ function from ‘lme4’ package^4, 8^ including species as a single random factor (intercept; Table S1B). The results of the two models resulted in virtually identical estimates for the fixed effects, so in the subsequent analyses we specified all models only within the ‘lmer’ framework, while also fitting study site as random intercept to control for non-independence of data points (to avoid problems of pseudo-replication arising from using multiple data points collected from the same study site).

We then attempted to replicate Kubelka et al.’s tests (their Figure 2AB and Table S2), while explicitly testing the evidence for differences in predation rates across geographic zones (i.e. using interactions). We thus fitted ‘mean year’ (quadratic) in interaction with ‘geographical zone’ (five-level factor). We then compared this model with three simpler models (Table S2, S4, Table 1): first, identical to the previous model but without the interaction; second model with the linear term ‘mean year’ in interaction with ‘geographical zone’, and a third model without this interaction (i.e. models we expected to find, but did not find, in Kubelka et al.’s Table S2). As the presumed increase in the Arctic predation rates (Figure 2AB^3^) occurred only after the year 2000, we also used the best fitting of the two interaction models (Table 1, Table S4) on data limited to after the year 1999 (Table S5A, *N* = 94 populations).

We found that predation rates were similar across geographical zones, except for the Southern Temperate zone, which had lower predation rates than the other zones (Figure 1AB, Table S2). Overall, the temporal change in predation rates was also similar across geographical zones (Figure 1AB, Table S2), even if we limit the data to the period after year 1999 when the change - according to Kubelka et al. - should have occurred (Table S5A). Importantly, the models without interaction were about 18 to 34 times more likely to be supported by the data than models with the interaction (Table 1 and S4).

#### Latitude

Using Kubelka et al.’s model (see their Table S6A), we first tested how patterns of predation rates changed over latitude by including a three-way interaction between ‘hemisphere’ (Northern or Southern), ‘mean year’ and ‘absolute latitude’ (Table S1C). We then also specified a similar model but using ‘lmer’ and species as a single random factor (intercept; Table S1D). The results of the two models were also identical, so in the subsequent analyses we specify all models only within ‘lmer’ framework, while fitting also study site as random intercept to to account for non-independence of data collected in the same study site.

We then attempted to replicate the Kubelka et al.’s tests (from their Figure 3AB and Table S6), while explicitly testing whether temporal trends in predation rates varied with latitude (i.e. using interactions). We thus fitted (Table S3) one model with ‘latitude’ (third-order polynomial) in interaction with ‘mean year’ of the study; second model with three-way interaction of ‘hemisphere’ (Southern or Northern), ‘absolute latitude’ and ‘mean year’; third model with ‘latitude’ (third-order polynomial) in interaction with ‘period’ (before or after year 2000); and fourth model with three-way interaction of ‘hemisphere’ (Southern or Northern), ‘absolute latitude’ and ‘period’ (before or after year 2000). We then compared these models to their simpler alternatives without any interactions (Table 2 and S4). Note that we have used a third-order polynomial of latitude to mimic the relationship Kubelka et al. depicted in their Fig.3.

In accordance with the results on geographical zones (Table S2), we found that predation rates were lower in the Southern hemisphere and increased globally over time, but without changing the latitudinal pattern (Table S3, S4 and 2). Importantly, the models without interactions were better supported by the data than models with interactions and models with ‘period’ (i.e. testing for the relationship presented by Kubelka et al.’s Figure 3) performed the worst of all models, receiving 60 to 130 times less empirical support than the best-supported models (Table 2 and S4).

#### Overall

Comparing the model for ‘Geographical zones’ together with the models for ‘Latitude’, we found that simple models without interactions fit the data better than models with interactions (Table 2 and S4).

**Table S1.** Predation rates in relation to mean year of the study and geography without controlling for study site.

| Model | Effect type | Response<br>Effect | ln(Daily predation rate + 0.01) |  |  | Total predation rate |  |  |
| --- | --- | --- | --- | --- | --- | --- | --- | --- |
|  |  |  | Estimate | 95%CI |  | Estimate | 95% CI |  |
| **A. Zone ‘Imekin’**<br>(Kubelka’s Table S2A<br>but with interaction) | Fixed | Intercept (Arctic) | -3.285 | -3.544 | -3.026 | 0.502 | 0.389 | 0.614 |
|  |  | ln (# of nests) | -0.007 | -0.070 | 0.056 | -0.001 | -0.028 | 0.026 |
|  |  | Mean year of the study | 0.274 | 0.160 | 0.389 | 0.111 | 0.063 | 0.16 |
|  |  | Zone - N. Temperate | -0.052 | -0.227 | 0.124 | -0.003 | -0.078 | 0.072 |
|  |  | Zone - N. Tropics | 0.102 | -0.199 | 0.404 | 0.055 | -0.076 | 0.185 |
|  |  | Zone - S. Temperate | -0.507 | -0.756 | -0.258 | -0.193 | -0.300 | -0.085 |
|  |  | Zone - S. Tropics | -0.179 | -0.493 | 0.136 | -0.045 | -0.179 | 0.089 |
|  |  | Mean year × N. Temperate | -0.076 | -0.231 | 0.08 | -0.016 | -0.082 | 0.05 |
|  |  | Mean year × N. Tropics | -0.195 | -0.489 | 0.099 | -0.084 | -0.209 | 0.042 |
|  |  | Mean year × S. Temperate | -0.154 | -0.435 | 0.127 | -0.069 | -0.188 | 0.051 |
|  |  | Mean year × S. Tropics | -0.136 | -0.432 | 0.159 | -0.048 | -0.173 | 0.077 |
|  | Random<br>(species) | Reciprocal of # of nests matrix | 8% |  |  | 5% |  |  |
|  |  | Phylogenetic matrix | 1% |  |  | 1% |  |  |
|  |  | Geographical distance matrix | 0% |  |  | 0% |  |  |
|  |  | Residual variance | 91% |  |  | 94% |  |  |
| **B. Zone ‘Imer’** | Fixed | Intercept (Arctic) | -3.247 | -3.513 | -2.984 | 0.522 | 0.410 | 0.637 |
|  |  | ln (# of nests) | -0.017 | -0.082 | 0.049 | -0.006 | -0.034 | 0.022 |
|  |  | Mean year of the study | 0.274 | 0.154 | 0.393 | 0.110 | 0.060 | 0.163 |
|  |  | Zone - N. Temperate | -0.040 | -0.225 | 0.145 | 0.003 | -0.079 | 0.084 |
|  |  | Zone - N. Tropics | 0.115 | -0.195 | 0.427 | 0.066 | -0.067 | 0.202 |
|  |  | Zone - S. Temperate | -0.507 | -0.760 | -0.239 | -0.191 | -0.302 | -0.081 |
|  |  | Zone - S. Tropics | -0.146 | -0.479 | 0.185 | -0.026 | -0.173 | 0.118 |
|  |  | Mean year × N. Temperate | -0.077 | -0.245 | 0.086 | -0.017 | -0.088 | 0.053 |
|  |  | Mean year × N. Tropics | -0.202 | -0.508 | 0.101 | -0.088 | -0.219 | 0.039 |
|  |  | Mean year × S. Temperate | -0.161 | -0.464 | 0.128 | -0.073 | -0.193 | 0.049 |
|  |  | Mean year × S. Tropics | -0.131 | -0.43 | 0.181 | -0.039 | -0.167 | 0.091 |
|  | Random | Species (intercept) | 10% |  |  | 13% |  |  |
|  |  | Residual variance | 90% |  |  | 87% |  |  |
| **C. Latitude ‘Imekin’**<br>(Kubelka’s Table S6A<br>but with interaction) | Fixed | Intercept (Northern) | -3.263 | -3.525 | -3.001 | 0.517 | 0.402 | 0.632 |
|  |  | ln (# of nests) | -0.017 | -0.076 | 0.042 | -0.004 | -0.029 | 0.021 |
|  |  | Hemisphere (Southern) | -0.662 | -1.005 | -0.319 | -0.271 | -0.418 | -0.125 |
|  |  | Mean Year of the study | 0.218 | 0.144 | 0.291 | 0.094 | 0.063 | 0.125 |
|  |  | Latitude (absolute) | -0.014 | -0.100 | 0.072 | -0.010 | -0.048 | 0.028 |
|  |  | Year × Hemisphere | -0.229 | -0.628 | 0.170 | -0.114 | -0.283 | 0.054 |
|  |  | Latitude × Hemisphere | -0.256 | -0.539 | 0.027 | -0.109 | -0.230 | 0.011 |
|  |  | Year × Latitude | 0.072 | -0.007 | 0.152 | 0.027 | -0.007 | 0.061 |
|  |  | Year × Latitude x Hemisphere | -0.181 | -0.489 | 0.126 | -0.084 | -0.213 | 0.046 |
|  | Random<br>(species) | Reciprocal of # of nests matrix | 11% |  |  | 6% |  |  |
|  |  | Phylogenetic matrix | 0% |  |  | 0% |  |  |
|  |  | Geographical distance matrix | 0% |  |  | 0% |  |  |
|  |  | Residual variance | 89% |  |  | 93% |  |  |
| **D. Latitude ‘Imer’** | Fixed | Intercept (Northern) | -3.21 | -3.482 | -2.943 | 0.547 | 0.429 | 0.665 |
|  |  | ln (# of nests) | -0.028 | -0.091 | 0.035 | -0.010 | -0.037 | 0.017 |
|  |  | Hemisphere (Southern) | -0.686 | -1.033 | -0.342 | -0.283 | -0.432 | -0.137 |
|  |  | Mean Year of the study | 0.210 | 0.133 | 0.288 | 0.090 | 0.058 | 0.123 |
|  |  | Latitude (absolute) | -0.015 | -0.105 | 0.073 | -0.015 | -0.054 | 0.025 |
|  |  | Year × Hemisphere | -0.255 | -0.657 | 0.147 | -0.125 | -0.292 | 0.046 |
|  |  | Latitude × Hemisphere | -0.279 | -0.566 | 0.017 | -0.117 | -0.242 | 0.004 |
|  |  | Year × Latitude | 0.075 | -0.006 | 0.158 | 0.030 | -0.006 | 0.065 |
|  |  | Year × Latitude x Hemisphere | -0.21 | -0.526 | 0.112 | -0.099 | -0.233 | 0.034 |
|  | Random | Species (intercept) | 12% |  |  | 15% |  |  |
|  |  | Residual variance | 88% |  |  | 85% |  |  |

Shown are model estimates and 95% confidence intervals (CI) and random variances calculated from ‘Imekin’ model output<sup>2</sup> (**A, C**) and the posterior estimates (medians) of the effect sizes with the 95% credible intervals (CI) from a posterior distribution of 5,000 simulated values generated by the ‘sim’ function in R<sup>6</sup> (**B, D**). Variance components were estimated by the ‘Imer’ function in R<sup>4</sup>. Mean year and absolute latitude were z-transformed (by subtracting the mean and dividing by standard deviation).
N = 237 populations representing 111 species.

**Table S2.** Predation rates in relation to mean year of the study and geographical zone, controlling for study site.

| Model | Effect type | Response<br>Effect | ln(Daily predation rate + 0.01) |  |  | Total predation rate |  |  |
| --- | --- | --- | --- | --- | --- | --- | --- | --- |
|  |  |  | Estimate | 95%CI |  | Estimate | 95% CI |  |
| **A. Simple & linear**<br>(Year + Zone) | Fixed | Intercept (Arctic) | -3.468 | -3.736 | -3.200 | 0.435 | 0.319 | 0.548 |
|  |  | ln (# of nests) | 0.047 | -0.009 | 0.104 | 0.021 | -0.003 | 0.045 |
|  |  | Mean year of the study | 0.147 | 0.064 | 0.225 | 0.065 | 0.032 | 0.098 |
|  |  | Zone - N. Temperate | -0.085 | -0.304 | 0.132 | -0.016 | -0.112 | 0.080 |
|  |  | Zone - N. Tropics | 0.069 | -0.250 | 0.395 | 0.044 | -0.090 | 0.186 |
|  |  | Zone - S. Temperate | -0.549 | -0.853 | -0.261 | -0.213 | -0.338 | -0.087 |
|  |  | Zone - S. Tropics | -0.219 | -0.569 | 0.130 | -0.061 | -0.212 | 0.089 |
|  | Random | Study site (intercept) | 61% |  |  | 62% |  |  |
|  |  | Species (intercept) | 2% |  |  | 3% |  |  |
|  |  | Residual variance | 36% |  |  | 36% |  |  |
| **B. Interaction & linear**<br>(Year × Zone) | Fixed | Intercept (Arctic) | -3.484 | -3.750 | -3.214 | 0.429 | 0.312 | 0.547 |
|  |  | ln (# of nests) | 0.044 | -0.014 | 0.102 | 0.020 | -0.004 | 0.045 |
|  |  | Mean year of the study | 0.242 | 0.078 | 0.398 | 0.092 | 0.021 | 0.161 |
|  |  | Zone - N. Temperate | -0.059 | -0.286 | 0.169 | -0.008 | -0.104 | 0.09 |
|  |  | Zone - N. Tropics | 0.102 | -0.237 | 0.457 | 0.059 | -0.091 | 0.203 |
|  |  | Zone - S. Temperate | -0.514 | -0.826 | -0.203 | -0.198 | -0.329 | -0.067 |
|  |  | Zone - S. Tropics | -0.176 | -0.550 | 0.180 | -0.049 | -0.209 | 0.104 |
|  |  | Mean year × N. Temperate | -0.109 | -0.302 | 0.085 | -0.026 | -0.110 | 0.059 |
|  |  | Mean year × N. Tropics | -0.138 | -0.468 | 0.180 | -0.049 | -0.190 | 0.092 |
|  |  | Mean year × S. Temperate | -0.219 | -0.538 | 0.107 | -0.095 | -0.232 | 0.043 |
|  |  | Mean year × S. Tropics | -0.138 | -0.498 | 0.212 | -0.034 | -0.186 | 0.116 |
|  | Random | Study site (intercept) | 63% |  |  | 63% |  |  |
|  |  | Species (intercept) | 2% |  |  | 3% |  |  |
|  |  | Residual variance | 35% |  |  | 34% |  |  |
| **C. Simple & quadratic**<br>(Year (quadratic) + Zone) | Fixed | Intercept (Arctic) | -3.454 | -3.724 | -3.174 | 0.443 | 0.329 | 0.557 |
|  |  | ln (# of nests) | 0.043 | -0.016 | 0.101 | 0.019 | -0.005 | 0.043 |
|  |  | Mean year (1<sup>st</sup> polynomial) | 2.212 | 0.975 | 3.401 | 0.965 | 0.459 | 1.469 |
|  |  | Mean year (2<sup>nd</sup> polynomial) | -0.600 | -1.696 | 0.514 | -0.384 | -0.870 | 0.095 |
|  |  | Zone - N. Temperate | -0.081 | -0.291 | 0.136 | -0.015 | -0.108 | 0.083 |
|  |  | Zone - N. Tropics | 0.073 | -0.249 | 0.400 | 0.049 | -0.087 | 0.184 |
|  |  | Zone - S. Temperate | -0.556 | -0.843 | -0.258 | -0.217 | -0.348 | -0.089 |
|  |  | Zone - S. Tropics | -0.215 | -0.563 | 0.144 | -0.06 | -0.209 | 0.091 |
|  | Random | Study site (intercept) | 62% |  |  | 63% |  |  |
|  |  | Species (intercept) | 2% |  |  | 2% |  |  |
|  |  | Residual variance | 36% |  |  | 35% |  |  |
| **D. Interaction & quadratic**<br>(Year(quadratic) × Zone) | Fixed | Intercept (Arctic) | -3.462 | -3.743 | -3.190 | 0.440 | 0.322 | 0.560 |
|  |  | ln (# of nests) | 0.046 | -0.010 | 0.106 | 0.020 | -0.005 | 0.045 |
|  |  | Mean year (1<sup>st</sup> polynomial) | 1.779 | -1.666 | 5.137 | 0.787 | -0.670 | 2.260 |
|  |  | Mean year (2<sup>nd</sup> polynomial) | 3.250 | -0.587 | 7.081 | 1.045 | -0.554 | 2.666 |
|  |  | Zone - N. Temperate | -0.096 | -0.329 | 0.134 | -0.017 | -0.113 | 0.083 |
|  |  | Zone - N. Tropics | 0.088 | -0.259 | 0.429 | 0.052 | -0.092 | 0.200 |
|  |  | Zone - S. Temperate | -0.575 | -0.905 | -0.256 | -0.224 | -0.360 | -0.090 |
|  |  | Zone - S. Tropics | -0.177 | -0.553 | 0.193 | -0.049 | -0.205 | 0.110 |
|  |  | Year (1<sup>st</sup> poly) × N. Temperate | -0.501 | -4.455 | 3.504 | -0.131 | -1.799 | 1.542 |
|  |  | Year (2<sup>nd</sup> poly) × N. Temperate | -4.614 | -8.780 | -0.491 | -1.675 | -3.445 | 0.040 |
|  |  | Year (1<sup>st</sup> poly) × N. Tropics | -0.769 | -7.280 | 5.586 | -0.244 | -2.887 | 2.558 |
|  |  | Year (2<sup>nd</sup> poly) × N. Tropics | -1.629 | -10.729 | 7.659 | -0.659 | -4.500 | 2.925 |
|  |  | Year (1<sup>st</sup> poly) × S. Temperate | -0.428 | -6.523 | 5.408 | -0.428 | -2.988 | 2.163 |
|  |  | Year (2<sup>nd</sup> poly) × S. Temperate | -5.683 | -12.527 | 1.191 | -2.143 | -5.126 | 0.777 |
|  |  | Year (1<sup>st</sup> poly) × S. Tropics | 0.385 | -5.306 | 6.282 | 0.321 | -2.273 | 2.806 |
|  |  | Year (2<sup>nd</sup> poly) × S. Tropics | -8.370 | -15.921 | -1.033 | -3.294 | -6.445 | -0.188 |
|  | Random | Study site (intercept) | 66% |  |  | 66% |  |  |
|  |  | Species (intercept) | 1% |  |  | 2% |  |  |
|  |  | Residual variance | 33% |  |  | 32% |  |  |

Shown are the posterior estimates (medians) of the effect sizes with the 95% credible intervals from a posterior distribution of 5,000 simulated values generated by the 'sim' function in R<sup>6</sup>. Variance components were estimated by the 'lmer' function in R<sup>4</sup>. Unless quadratics, mean year was z-transformed (by subtracting the mean and dividing by standard deviation).
N = 237 populations representing 111 species.

**Table S3year.**
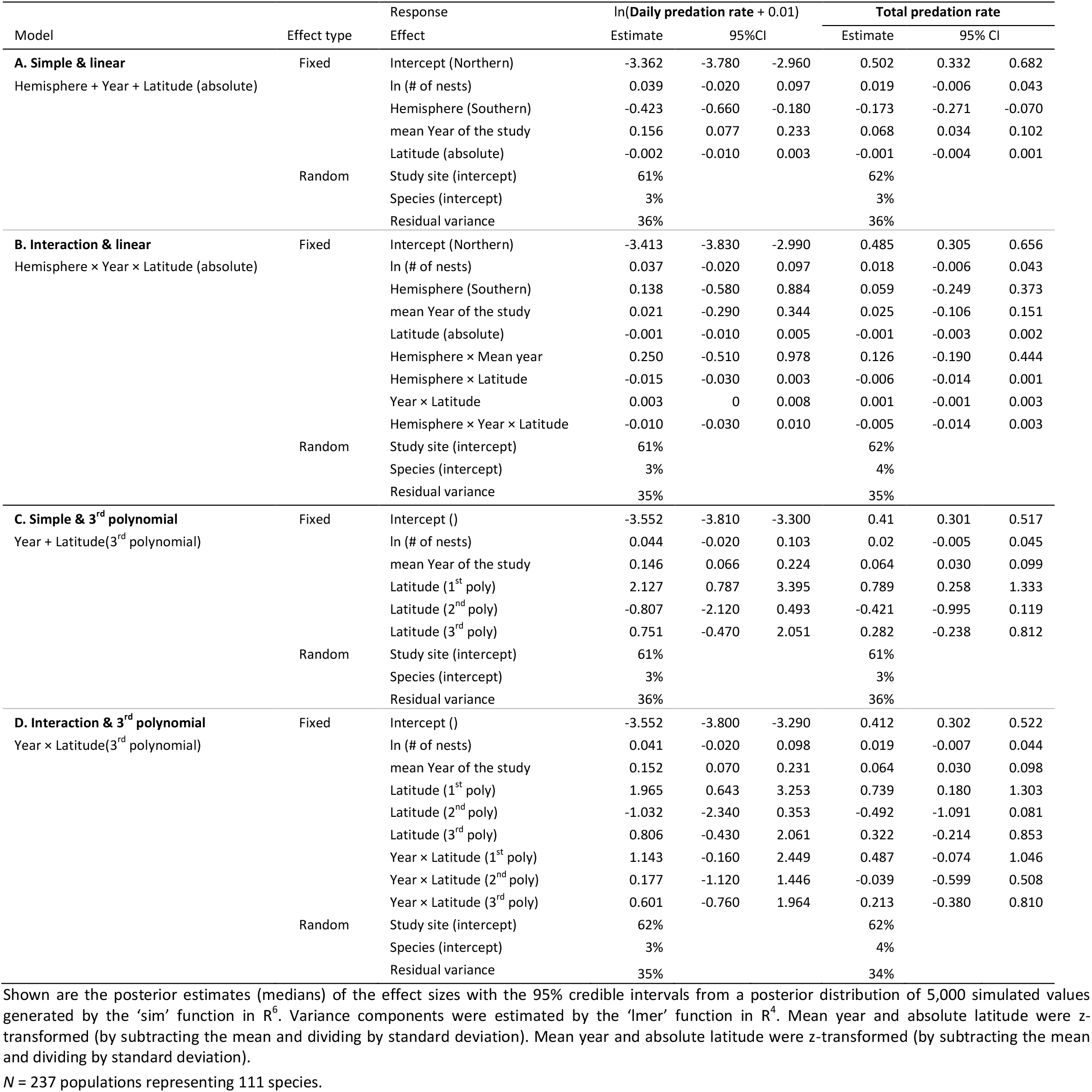
Predation rates in relation to mean year and latitude of the study, controlling for study site and year.

**Table S3period.** Predation rates in relation to mean year and latitude of the study, controlling for study site and year.

| Model | Effect type | Response | ln(Daily predation rate + 0.01) |  |  | Total predation rate |  |  |
| --- | --- | --- | --- | --- | --- | --- | --- | --- |
|  |  | Effect | Estimate | 95%CI |  | Estimate | 95% CI |  |
| **E. Simple & linear**<br>Hemisphere + Period +<br>Latitude (absolute) | Fixed | Intercept (Northern & after 2000) | -3.307 | -3.740 | -2.890 | 0.529 | 0.346 | 0.709 |
|  |  | ln (# of nests) | 0.044 | -0.010 | 0.103 | 0.020 | -0.005 | 0.045 |
|  |  | Hemisphere (Southern) | -0.391 | -0.640 | -0.150 | -0.162 | -0.267 | -0.052 |
|  |  | Period (before 2000) | -0.176 | -0.340 | -0.010 | -0.065 | -0.135 | 0.003 |
|  |  | Latitude (absolute) | -0.002 | -0.010 | 0.004 | -0.001 | -0.004 | 0.001 |
|  | Random | Study site (intercept) | 62% |  |  | 62% |  |  |
|  |  | Species (intercept) | 5% |  |  | 7% |  |  |
|  |  | Residual variance | 33% |  |  | 31% |  |  |
| **F. Interaction & linear**<br>Hemisphere × period ×<br>Latitude (absolute) | Fixed | Intercept (Northern & after 2000) | -3.673 | -4.210 | -3.130 | 0.380 | 0.148 | 0.612 |
|  |  | ln (# of nests) | 0.047 | -0.010 | 0.105 | 0.020 | -0.006 | 0.045 |
|  |  | Hemisphere (Southern) | 0.592 | -0.450 | 1.651 | 0.302 | -0.154 | 0.770 |
|  |  | Period (before 2000) | 0.364 | -0.260 | 0.999 | 0.165 | -0.114 | 0.429 |
|  |  | Latitude (absolute) | 0.005 | -0 | 0.013 | 0.002 | -0.002 | 0.005 |
|  |  | Hemisphere × Period | -0.647 | -2.130 | 0.798 | -0.352 | -0.980 | 0.277 |
|  |  | Hemisphere × Latitude | -0.030 | -0.060 | 0 | -0.014 | -0.027 | -0.001 |
|  |  | Period × Latitude | -0.011 | -0.020 | 0 | -0.005 | -0.009 | 0 |
|  |  | Hemisphere × Period × Latitude | 0.022 | -0.020 | 0.062 | 0.012 | -0.005 | 0.028 |
|  | Random | Study site (intercept) | 62% |  |  | 62% |  |  |
|  |  | Species (intercept) | 6% |  |  | 8% |  |  |
|  |  | Residual variance | 32% |  |  | 30% |  |  |
| **G. Simple & 3<sup>rd</sup> polynomial**<br>Hemisphere + Year +<br>Latitude(3<sup>rd</sup> polynomial) | Fixed | Intercept (Northern) | -3.478 | -3.750 | -3.220 | 0.438 | 0.324 | 0.554 |
|  |  | ln (# of nests) | 0.049 | -0.010 | 0.108 | 0.022 | -0.004 | 0.047 |
|  |  | Period (before 2000) | -0.159 | -0.330 | 0.004 | -0.059 | -0.129 | 0.009 |
|  |  | Latitude ( 1<sup>st</sup> poly) | 2.044 | 0.727 | 3.370 | 0.753 | 0.162 | 1.319 |
|  |  | Latitude ( 2<sup>nd</sup> poly) | -0.634 | -1.980 | 0.748 | -0.369 | -0.956 | 0.229 |
|  |  | Latitude ( 3<sup>rd</sup> poly) | 0.918 | -0.390 | 2.202 | 0.370 | -0.169 | 0.913 |
|  | Random | Study site (intercept) | 62% |  |  | 62% |  |  |
|  |  | Species (intercept) | 5% |  |  | 7% |  |  |
|  |  | Residual variance | 33% |  |  | 31% |  |  |
| **H. Interaction & 3<sup>rd</sup> polynomial**<br>Hemisphere × Period ×<br>Latitude(3<sup>rd</sup> polynomial) | Fixed | Intercept ( ) | -3.472 | -3.750 | -3.210 | 0.443 | 0.333 | 0.559 |
|  |  | ln (# of nests) | 0.048 | -0.010 | 0.108 | 0.020 | -0.005 | 0.045 |
|  |  | Period (before 2000) | -0.178 | -0.340 | -0.010 | -0.067 | -0.141 | 0.004 |
|  |  | Latitude (1<sup>st</sup> poly) | 3.677 | 1.692 | 5.653 | 1.499 | 0.681 | 2.347 |
|  |  | Latitude (2<sup>nd</sup> poly) | -0.197 | -2.110 | 1.702 | -0.215 | -1.059 | 0.597 |
|  |  | Latitude (3<sup>rd</sup> poly) | 1.962 | -0.200 | 4.085 | 0.878 | -0.070 | 1.798 |
|  |  | Year × Latitude (1<sup>st</sup> poly) | -2.874 | -5.300 | -0.440 | -1.291 | -2.304 | -0.262 |
|  |  | Year × Latitude (2<sup>nd</sup> poly) | -1.398 | -3.910 | 1.162 | -0.539 | -1.650 | 0.559 |
|  |  | Year × Latitude (3<sup>rd</sup> poly) | -1.522 | -4.120 | 1.107 | -0.713 | -1.792 | 0.437 |
|  | Random | Study site (intercept) | 62% |  |  | 62% |  |  |
|  |  | Species (intercept) | 6% |  |  | 8% |  |  |
|  |  | Residual variance | 32% |  |  | 30% |  |  |

**Table S4.** Model comparison for total nest predation rate.

| Model<sup>a</sup> | Predictors | # of parameters<sup>b</sup> | AIC | $\Delta AIC^c$ | $w_i^d$ | Cumulative $w_i^e$ | ER<sup>f</sup> |
| --- | --- | --- | --- | --- | --- | --- | --- |
| 1 | Year + Hemisphere + Latitude (absolute) | 5 | -53.50 | 0.00 | 0.28 | 0.28 | 1.00 |
| 2 | Year + Latitude (3rd polynomial) | 6 | -52.99 | 0.51 | 0.21 | 0.49 | 1.29 |
| 3 | Year (quadratic) + Geographical area | 8 | -52.92 | 0.58 | 0.21 | 0.70 | 1.34 |
| 4 | Year + Geographical area | 7 | -52.42 | 1.08 | 0.16 | 0.86 | 1.71 |
| 5 | Year × Hemisphere × Latitude (absolute) | 9 | -50.86 | 2.64 | 0.07 | 0.93 | 3.75 |
| 6 | Year × Latitude (3rd polynomial) | 9 | -50.00 | 3.50 | 0.05 | 0.98 | 5.76 |
| 7 | Year × Geographical area | 11 | -46.40 | 7.10 | 0.01 | 0.99 | 34.87 |
| 8 | Year (quadratic) × Geographical area | 16 | -45.76 | 7.74 | 0.01 | 0.99 | 47.90 |
| 9 | Period × Latitude (3rd polynomial) | 9 | -43.88 | 9.62 | 0 | 1 | 122.81 |
| 10 | Period × Hemisphere × Latitude (absolute) | 9 | -43.28 | 10.22 | 0 | 1 | 165.44 |
| 11 | Period + Hemisphere + Latitude (absolute) | 5 | -41.87 | 11.63 | 0 | 1 | 334.46 |
| 12 | Period + Latitude (3rd polynomial) | 6 | -40.27 | 13.23 | 0 | 1 | 746.23 |

<sup>a</sup>Each model is fitted with maximum likelihood and controlled for number of nests in a given population (ln-transformed) and for multiple populations at given site or for a given species using site and species as random intercepts (i.e. all models have same random structure). Predictors are Year (mean year of the study), Hemisphere (Northern vs Southern), Latitude (degrees), Geographical area (Arctic, North temperate, North tropics, South tropics, South temperate,,), and Period (historic: 1944-1999 vs. recent: 2000-2016). Models that include Period (instead of Year) are not supported by the data (69-320 times less likely than the best model). Models including the interaction between time and geographical/latitude do not improve the model fit or are much less supported by the data than models without the interaction. For model outputs see Table S2-3.
<sup>b</sup>Number of model parameters without the random effects. <sup>c</sup>The difference in AICc between the first-ranked model and the given model.
<sup>d</sup>Akaike weight – the weight of evidence that a given model is the best approximating model (i.e., probability of the model).
<sup>e</sup>Cumulative Akaike weight, <sup>f</sup>Evidence ratio – model weight of the first-ranked model relative to that of the given model (i.e., how many times is the first-ranked model more likely than the given model).

**Table S5.** Predation rates in relation to mean year of the study and geographical zone for limited datasets.

| Model | Effect type | Response<br>Effect | ln(Daily predation rate + 0.01) |  |  | Total predation rate |  |  |
| --- | --- | --- | --- | --- | --- | --- | --- | --- |
|  |  |  | Estimate | 95%CI |  | Estimate | 95% CI |  |
| **A. Interaction & linear**<br>(Year × Zone)<br>Data with year >1999<br>N = 94 populations | Fixed | Intercept (Arctic) | -3.254 | -3.649 | -2.867 | 0.541 | 0.383 | 0.697 |
|  |  | ln (# of nests) | 0.041 | -0.038 | 0.120 | 0.016 | -0.015 | 0.048 |
|  |  | Mean year of the study | 0.199 | 0.026 | 0.378 | 0.060 | -0.010 | 0.129 |
|  |  | Zone - N. Temperate | -0.170 | -0.520 | 0.184 | -0.054 | -0.206 | 0.095 |
|  |  | Zone - N. Tropics | -0.081 | -0.516 | 0.341 | -0.035 | -0.214 | 0.153 |
|  |  | Zone - S. Temperate | -0.644 | -1.081 | -0.193 | -0.276 | -0.468 | -0.078 |
|  |  | Zone - S. Tropics | -0.278 | -0.770 | 0.211 | -0.08 | -0.295 | 0.132 |
| | | Mean year × N. Temperate | -0.102 | -0.410 | 0.212 | -0.011 | -0.139 | 0.118 |
| | | Mean year × N. Tropics | 0.057 | -0.320 | 0.422 | 0.046 | -0.112 | 0.198 |
| | | Mean year × S. Temperate | -0.122 | -0.493 | 0.255 | -0.019 | -0.175 | 0.139 |
| | | Mean year × S. Tropics | -0.598 | -1.209 | 0.009 | -0.217 | -0.479 | 0.043 |
|  | Random | Study site (intercept) | 73% |  |  | 79% |  |  |
|  |  | Species (intercept) | 0% |  |  | 3% |  |  |
|  |  | Residual variance | 27% |  |  | 18% |  |  |
| **B. Mean year > 1970**<br>Data with year >1970<br>N = 226 populations | Fixed | Intercept (Arctic) | -3.536 | -3.792 | -3.269 | 0.419 | 0.308 | 0.528 |
|  |  | ln (# of nests) | 0.044 | -0.015 | 0.104 | 0.021 | -0.004 | 0.045 |
|  |  | Mean year of the study | 0.071 | -0.017 | 0.159 | 0.029 | -0.009 | 0.067 |
|  | Random | Study site (intercept) | 65% |  |  | 66% |  |  |
|  |  | Species (intercept) | 3% |  |  | 3% |  |  |
|  |  | Residual variance | 32% |  |  | 32% |  |  |

Shown are the posterior estimates (medians) of the effect sizes with the 95% credible intervals from a posterior distribution of 5,000 simulated values generated by the 'sim' function in R<sup>6</sup>. Variance components were estimated by the 'lmer' function in R<sup>4</sup>. Mean year was z-transformed (by subtracting the mean and dividing by standard deviation).

### Exploring the temporal change in predation rates

The general increase in predation rates found by Kubelka et al. — and confirmed in our analyses — can arise if field protocols and/or statistical methods change over time. In Kubelka et al.’s dataset, 59% (total N = 237) of populations lack the number of exposure days (i.e. the total number of days that nests were followed from finding until the nest finished (hatched, depredated, failed to other causes) that are needed to calculate daily predation rates according to Mayfield^9^, the method used by the Kubelka et al.^3^. Kubelka et al. derive such exposure days using nesting period (egg-laying + incubation period) of the species and a conversion coefficient introduced by Beintema^10^, which indicates how much of the incubation period (in case of Kubelka et al. of the nesting period) was observed, i.e. indicating when the nests were generally found. Kubelka et al. assumed that 0.9 of nesting period was observed if nests were found close to laying or nests searched daily, 0.6 if nests were found early in the nesting period or nests searched once-twice a week, or 0.5 if nests were found in the middle of the nesting period (*N*_0.5_ = 11, *N*_0.6_ = 114, N_0.9_ = 14 populations). In other words, Kubelka et al. assumed that the vast majority of nests were found earlier than in the middle of the nesting period. However, such an assumption might be too optimistic for many populations. Even in a recent, intensive research scheme with multiple nest surveys per week by ~2-6-person teams at various Arctic sites, nests are rarely found at laying (mean across sites = 0.35 of nesting period, range: 0.22 – 0.49; *N* = 10,716 nests from 16 sites monitored after 2000; Figure S1; using open-access data from the Arctic Shorebird Demographics Network^11^). Importantly, the need to use ‘Beintema conversions’ might have changed over time. We have thus explored five ways how such ‘Beintema conversions’ affect the temporal change in predation rates. Note that one Arctic population was indicated as transformed in the Kubelka et al.’s dataset but lacked the actual transformation value. Nevertheless, its exposure was indicated in the Kubelka et al.’s dataset and present also in the original reference, i.e. this population should have been indicated as not transformed and we use it in the subsequent analyses as such.

First, we visualized how the number of populations that required a ‘Beintema conversion’ changed over time (Figure 1G and S2; using locally estimated scatterplot smoothing). We reveal a steady decline in the number of studies lacking exposure data, i.e. studies where Kubelka et al. used the Beintema conversion. The decline is particularly dramatic after 2000, which corresponds with Kubelka et al.’s distinction between before and after 2000 period, and especially in Arctic which corresponds with reported exponential increase in the predation rates in Arctic.

Second, we used the Kubelka et al.’s populations with known (i.e., termed “true” below) number of exposure days, known nesting period length, and known fates (N = 65) and estimated daily predation rates with varying conversion coefficients (0.5 × observed proportion of nesting period × nesting period × (number of nests depredated or failed to other causes) + (observed proportion of nesting period × nesting period × (number of hatched and infertile clutches). We then visualized the new daily predation rates against the original values to investigate how this method over- or under-estimates the daily predation rates. Despite the strong correlation between true daily predation rates (i.e. those extracted from the literature) and the newly derived ones^3^, we found severe over- and under-estimation depending on the ‘proportion of nesting period’ assumed for the calculations (Figure 1I and S3). If we assume that only 0.1-0.4 of the nesting period is observed, the predation rates are severely over-estimated for all (in case of 0.4 for most) original values (Figure 1I and SB). Assuming that nests are observed for half of the nesting period, overestimates the low true values and underestimates the larger ones. Assuming that nests are observed for longer than half of nesting period (>0.5), further overestimates the predation rates, including the lower true values.

Third, we explored how the increase in predation rates over time (Figure 1A-F) changes if we vary proportion of observed nesting period (i.e. Beintema’s coefficient) from 0.1 to 0.9 for populations with mean year <2000 and lacking exposure days (i.e. populations where Kubelka at al. used Beintema coefficient to calculate exposure). In other words, we assumed that intensive nest searching used by Kubelka (i.e. nests found before or during mid-nesting period) is always valid for data >2000, but uncertain for data <2000. To each dataset we fitted a model with ‘mean year’ of the study as a fixed effect, controlling for number of nest (ln-transformed) and site and species as random intercepts. We then plotted the model predictions (Figure 1H). This exercise revealed sensitivity of the data to the ‘Beintema conversion’ (Figure 1H) with conversion factors <0.5 (which were never used by Kubelka) generating statistically non-significant year effects, sometimes even in the opposite direction than reported by Kubelka et al.

Fourth, we tested for the effect of mean year on predation rates by using only data with known exposure days or predation rates (*N* = 98 populations; Table S6). First, we fitted two models: first with latitude (3^rd^ polynomial) in interaction with year, and second with three-way interaction of hemisphere, latitude (absolute) and year. Then, we fitted an additional two models using only Arctic (*N* = 46 populations) and North Temperate zone (*N* = 42) data (the other zones contained only 0-5 populations): first model with mean year (quadratic) in interaction with geographical zone, the second model with linear mean year in interaction with geographical zones. We then also fitted the same four models but without interactions (Table S6). We found no support for interactions, the geographical effect or the year effect (Table S6, Figure 1CF).

Fifth, we explored how the mean year effect changes when we exclude 10 sparsely distributed data points < 1970 (as all above mentioned models underestimate the effect of these populations). Using model with mean year as a predictor (same as Kubelka et al. in Table S2a) and site and species as random intercepts reduced the original Kubelka et al.’s year effect by 59% (Table S5B), revealing the influence of the 10 early data points.

**Figure S1.**
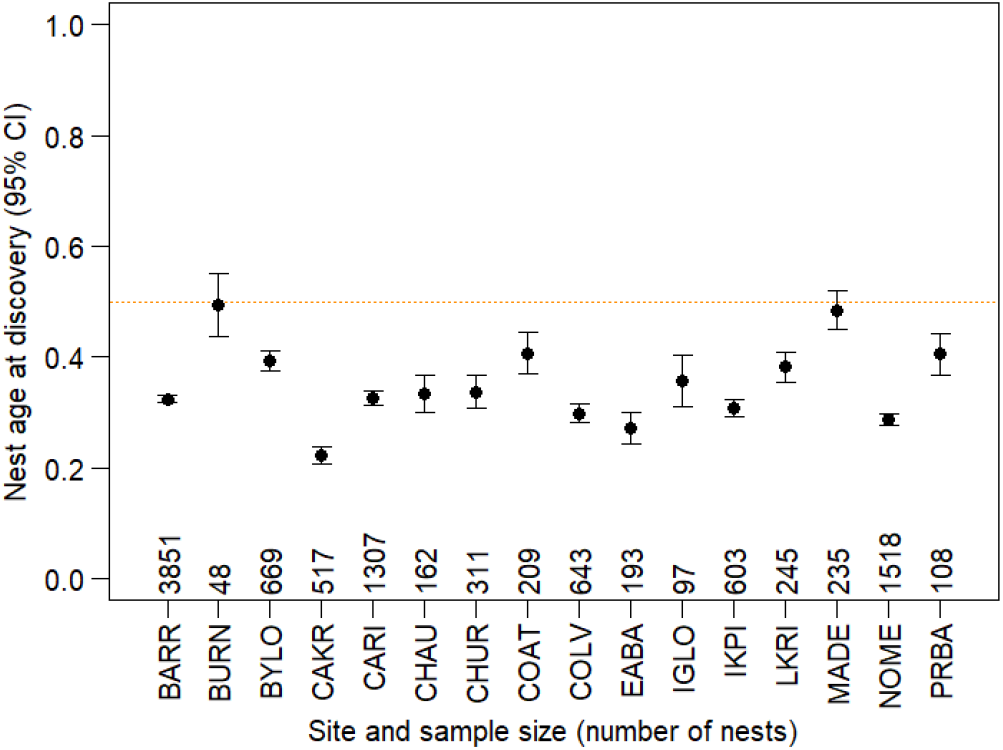
Nest age (proportion of the nesting period elapsed) at the time of nest discovery. Points indicate means, bars 95% Cis for each of 16 sites in the Arctic Shorebird Demographics Network in Russia, Alaska, and Canada (2003-2014). Numbers indicate number of nests. Horizontal dotted line indicates 0.5 (midpoint of the nesting period). For further information on these sites and nest-searching protocols see^11, 12^.

**Figure S2.**
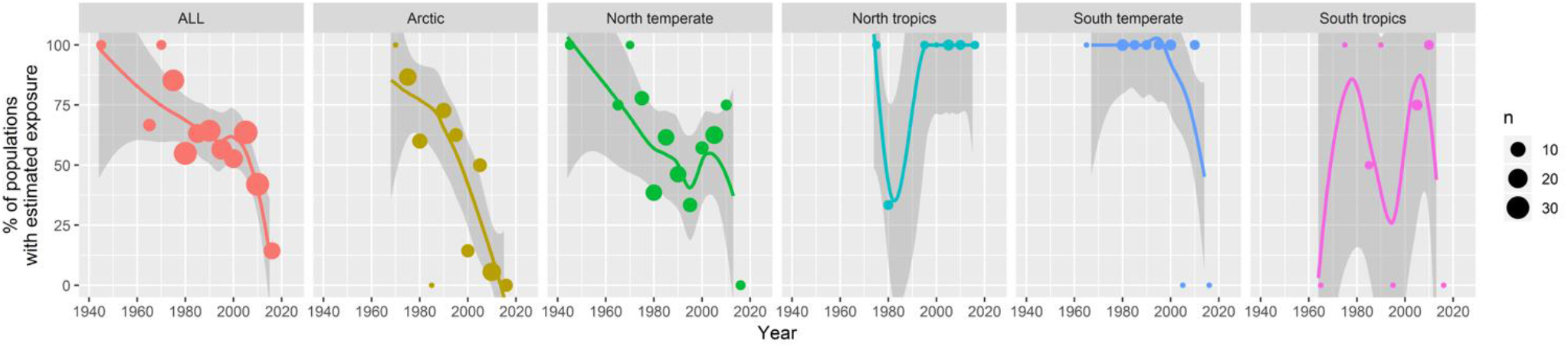
Temporal change in percentage of populations needing ‘Beintema conversion’ to estimate exposure. Dots represent percentages for 5-year intervals, lines and shaded areas locally estimated scatterplot smoothing with 95% confidence intervals

**Figure S3.**
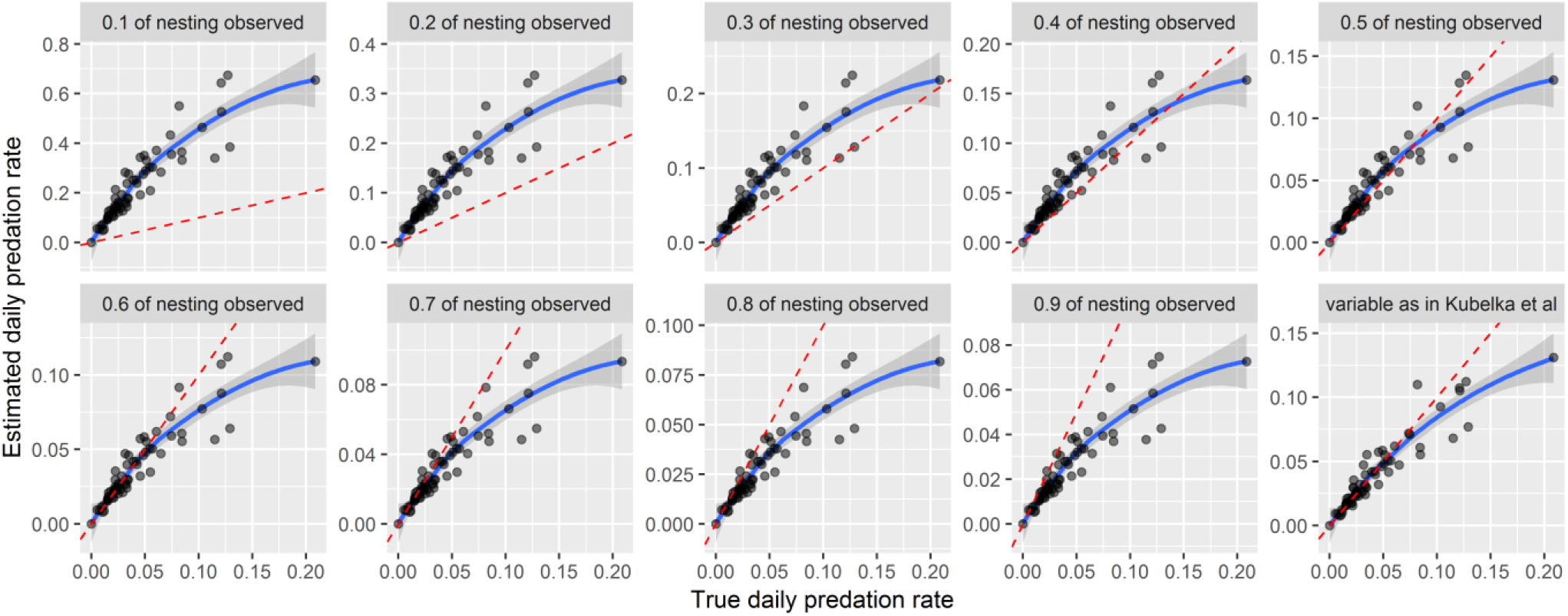
The assumption about the proportion of nesting period being observed influences daily predation rate estimation. Each dot represents one of 65 populations with true daily predation rates from the literature and all information needed to estimate daily predation rates for various proportions of the nesting period that is on average assumed to be observed (panel titles; note that the last panel uses proportions specific to each population as used by Kubelka et al.). Red dashed line indicates no difference between true values (x-axis) and estimated values (y-axis). Blue line with shaded area indicates locally estimated scatterplot smoothing with 95%CIs. Note that points and lines below the dashed lines indicates underestimation and above overestimation of the true values.

**Table S6a.**
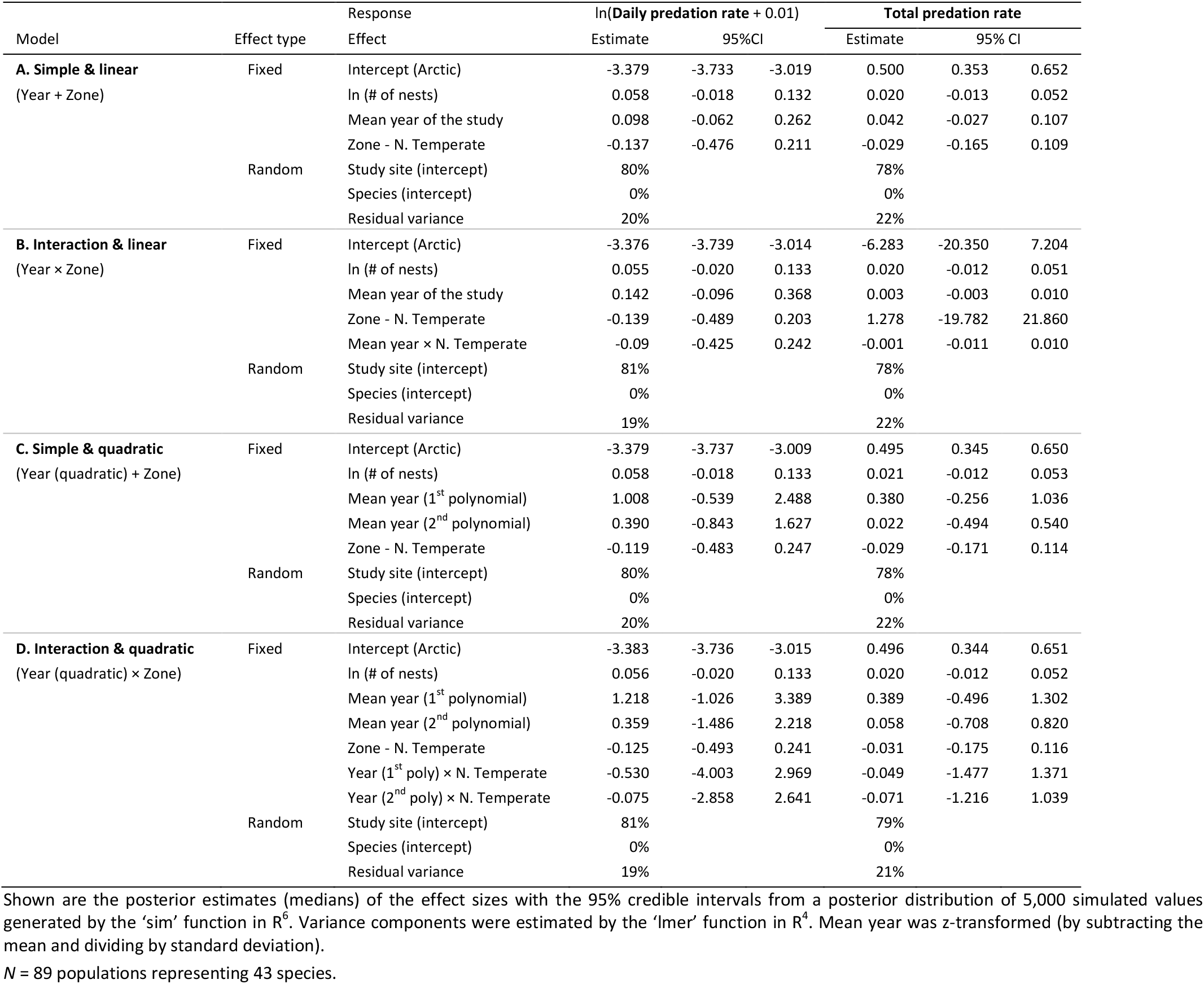
Predation rates in relation to mean year of the study and region (Arctic or N. temperate) using non-transformed data.

**Table S6b.**
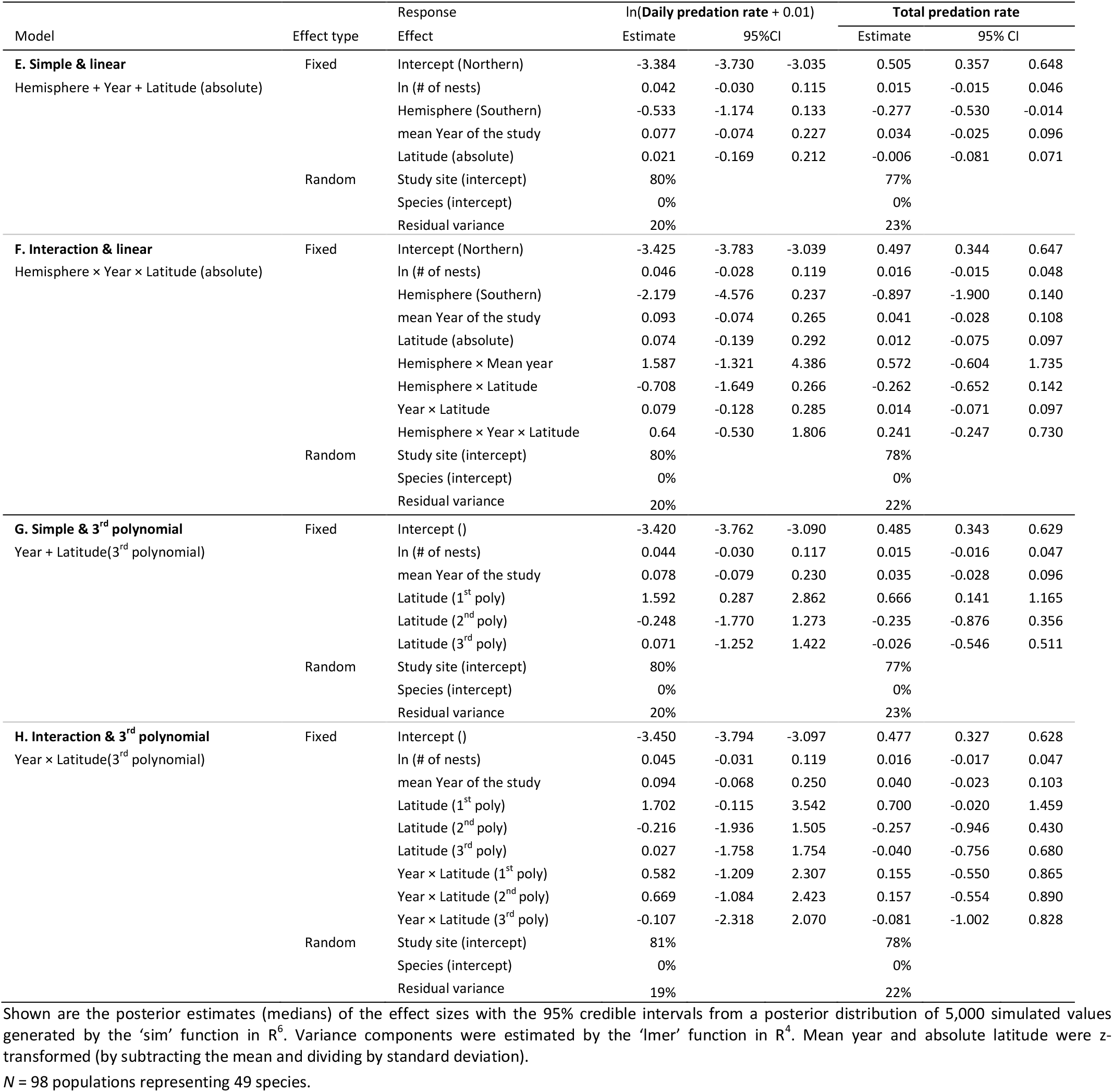
Predation rates in relation to mean year of the study and latitude using non-transformed data.

### Estimating repeatability of extracting information from the sources about ‘Beintema conversion’

For 38% of 128 populations (where Kubelka et al. assumed that more than 50% of nesting period was observed) we were unable to find information in the reference to suggest such assumpiton was appropriate. For sources where we found some relevant information about nest searching intensity and about when within nesting period most nests were found, a different person extracted the information a new for 73 sources. The conclusions differed in 30% of the sources.

### Exploring within-population changes in predation rates over time

Kubelka et al. tested for within-population change between periods (before and after 2000) in 9 populations at 7 sites and found a significant effect of period on the daily predation rates, where daily predation rates increased after 2000. We reviewed the references used by Kubelka et al. using their criteria for including populations (≥2 years and ≥12 nests with known fate for each period). We found information for a total of 23 populations. The 23 included 7 of the 9 included by Kubelka et al; for the remaining two, we were unable to obtain the necessary information for one (*Vanellus vanellus* in Czech Republic; Kubelka in litt.) and we found that the other population included only 13 nests after 2000 and the observation period was not known for most of those, so we excluded that population from further consideration *Calidris melanotos* at Kuparuk, Alaska^11^). One population not included by Kubelka et al. was from a low latitude (28° N); we excluded this population because, Kubelka et al. report the increased predation rates only for higher latitudes. For the remaining 22 populations (Table S7), we calculated daily predation rates based on the information we found in the literature or unpublished datasets, using the Beintema transformation when necessary (using 0.5 when we found no information to indicate that most nests were found prior to the midpoint of incubation, or 0.6 if nest-searching was conducted at least weekly or nest age at discovery was less than half of the nesting period). Our predation rate values occasionally differed from Kubelka et al.’s when we found additional data (years or nests) that were excluded by the Kubelka et al. or when we applied a different value for the Beintema transformation (Table S7).

We repeated Kubelka et al.’s assessment of within-population change in predation rates for our 22 populations by applying the same linear mixed-effects model, including fixed effects of period and latitude (scaled by subtracting the mean and dividing by standard deviation) and random effects of species and locality. Like Kubelka et al., we applied the model with package lme4 in R (Bates et al. 2014; R Core Team 2018). With our expanded dataset, we likewise found a positive effect of period (β_period_ = 0.29, 95% CI = 0.05 to 0.53, *p* = 0.03), indicating an increase in daily predation rates after 2000, although 46% smaller than the increase estimated by Kubelka et al. (β_period_ = 0.54, 95% CI = 0.11 to 0.97).

With the 22 populations, we then explored the consequences of the Beintema transformation for the apparent within-population change. We applied the above model separately to two groups: first, the populations for which the Beintema transformation was consistently needed (applied to both periods, or never applied; *N* = 13 populations at 5 sites; Figure S4a); and second, the populations that required the transformation in only one period, which was before 2000 in all cases (N = 9 populations at 3 sites; Figure S4b). For population with the consistent transformation, the effect of period dropped by 50% from our initial effect (β_period_ = 0.29) and became statistically non-significant (β_period_ = 0.14, 95% CI = −0.11 to 0.39, *p* = 0.28). For populations where the transformation was necessary only for the period before year 2000, the effect increased by 34% from our initial effect and remained significant (β_period_ = 0.49, SE = 0.20, *p* = 0.02). This suggests that using the Beintema transformation during only one of the two periods could explain the apparent effect of period on daily predation rates in the larger dataset.

Finally, for the 9 populations that required the transformation only before 2000, we conducted a sensitivity analysis for the value of the Beintema coefficient (*B*). Originally, we used *B* = 0.5 for all 9 populations because nest-searching was conducted less than weekly or no information was provided. However, as discussed above, at least in Arctic populations values higher than *B* = 0.6 (when nests are on average found just before the midpoint of the nesting period) are unlikely to be valid even in modern studies (see above), and *B* = 0.5 is sometimes more appropriate even with extensive nest-searching effort (Figure S1). Values lower than *B* = 0.5 were not considered by Kubelka et al., but would be appropriate if nests were found late in incubation or near hatching (Beintema 1996), which is likely for studies with less than weekly nest searching effort or for cryptic species. We thus varied Beintema coefficient from 0.1 to 0.4 to evaluate the sensitivity of the change in predation rate between periods to the assumptions made for the Beintema transformation. We then fitted the same model as above, using each value of *B* in turn. For this sensitivity analysis, we excluded one population for which the pre-2000 values were calculated from two different references, only one of which required the transformation (Whimbrel *Numenius phaeopus* at Churchill, Manitoba). We found that all values <0.5 resulted in a nonsignificant effect of period (p ≥ 0.14), and in the most extreme case (*B* = 0.1), the direction of the effect was opposite to the one found by Kubelka et al. and of the same magnitude (Figure S5, Table S8). In other words, smaller *B* values often produced higher daily predation estimates for before 2000 data than for after 2000 data (Figure S5), which often resulted in a conclusion that predation rate was not higher after 2000 than before 2000.

With no information provided in the sources for nest-searching frequency or age at which nests were found, it is impossible to tell which *B* value is most appropriate for many published studies. However, it seems likely that values of B < 0.5 would sometimes be appropriate for the studies from the 1960s and 1970s, especially if nests were found opportunistically or with low nest-searching effort. Given the sensitivity of the apparent change in daily predation rates to the value of *B* that was selected, and the lack of any change in daily predation rates in populations for which predation rates were known or *B* was applied consistently, the apparent increase in predation rates after 2000 detected by Kubelka et al. might have been a methodological artefact.

**Figure S4.**
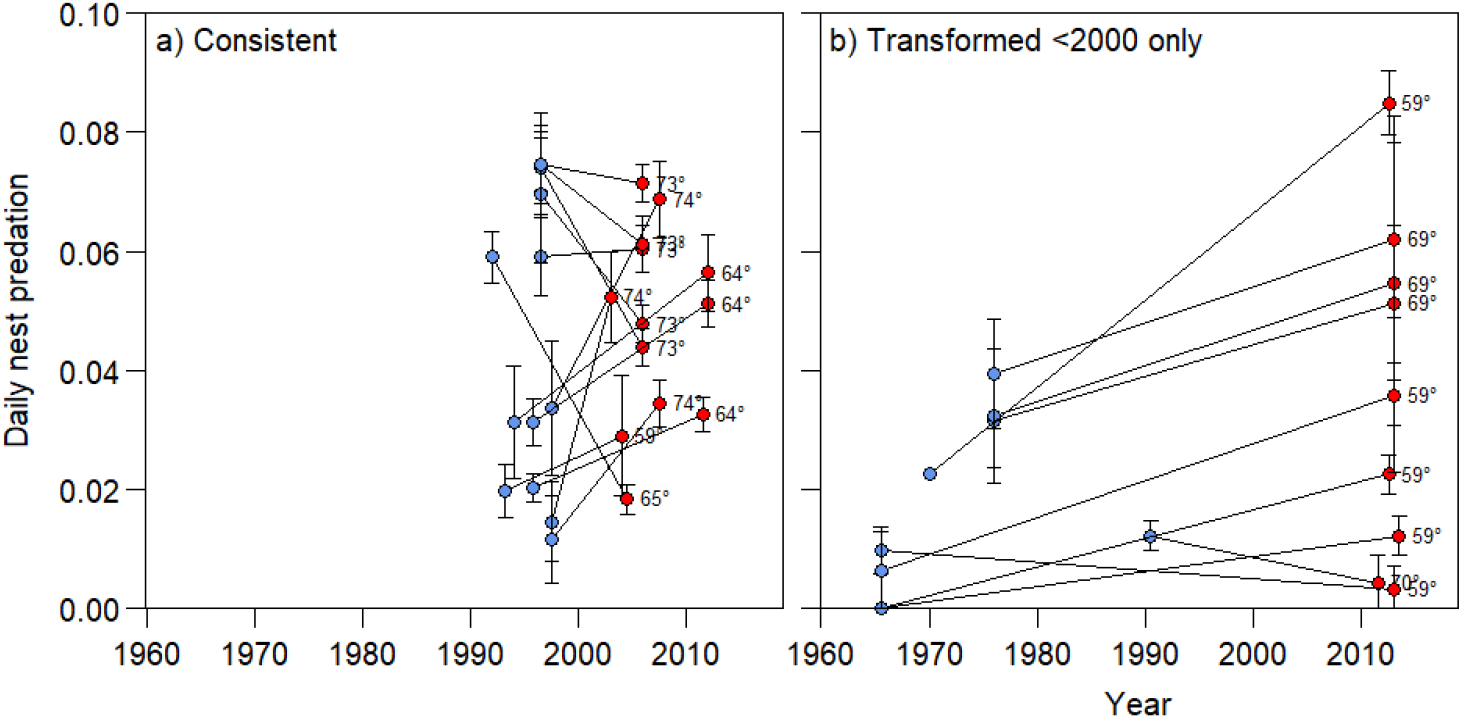
Population-specific change in nest predation over time. **a,b.** Populations that either consistently required the Beintema transformation in both periods, or consistently reported observation time explicitly (**a**), and populations that required the Beintema transformation in only one period (always before 2000; **b**). Points indicate means, bars 95% CIs. Colour indicates before 2000 (blue) and after 2000 (red), lines connect the same populations and numbers next to red points indicate the latitude of each population.

**Figure S5.**
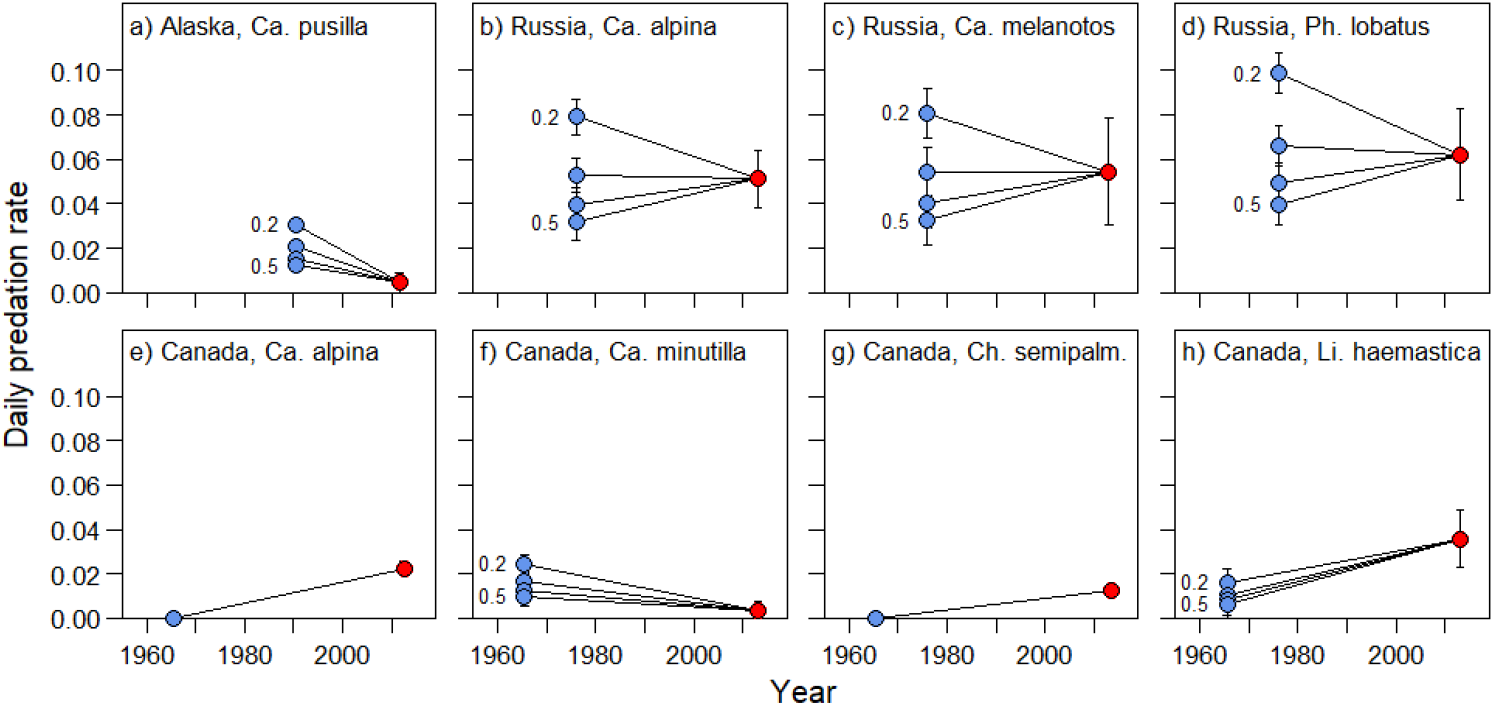
Population-specific daily predation rate according to species, location and conversion coefficient B. **a-h,** Each panel represents one population that required the Beintema transformation in only one period (always before 2000). Points indicate means, bars 95% CIs (calculated following^13^). Colour indicates before 2000 (blue) and after 2000 (red), numbers next to blue points indicate the various values of conversion coefficient (*B* = 0.2, 0.3, 0.4, or 0.5) used to estimated daily predation rate for before 2000 data. *B* = 0.1 was tested but often produced much higher predation rate values and is not shown for clarity. For two populations (**e, g**), predation rate before 2000 was always zero regardless of the conversion coefficient because zero nests were depredated. Details for each population are provided in Table S7.

**Table S7.** Shorebird populations used in re-analysis of within-population changes in daily predation rate from historic (<2000) to recent (≥2000) periods.

| Species<sup>a</sup> | Location | Latitude | Longitude | Period | DPR | SEM | N years | Mean year | N nests | Exposure | B<sup>b</sup> | Included by Kubelka et al.<sup>c</sup> | Source<sup>d</sup> |
| --- | --- | --- | --- | --- | --- | --- | --- | --- | --- | --- | --- | --- | --- |
| *Charadrius semipalmatus* | Canada | 58.698 | -93.942 | historic | 0 | 0 | 4 | 1966 | 15 | 196.0 | 0.5 | - | 1 |
|  |  |  |  | recent | 0.012 | 0.003 | 2 | 2014 | 67 | 1003.0 | - | - | 2 |
| *Limosa haemastica* | Canada | 58.698 | -93.942 | historic | 0.006 | 0.006 | 4 | 1966 | 12 | 155.2 | 0.5 | Yes | 1 |
|  |  |  |  | recent | 0.036 | 0.013 | 3 | 2013 | 20 | 201.5 | - | Yes | 2 |
| *Numenius phaeopus* | Canada | 58.698 | -93.942 | historic | 0.023 | NA | 16 | 1970 | 90 | 1121.0 | 0.5 | Yes<sup>1</sup> | 1, 3 |
|  |  |  |  | recent | 0.085 | 0.005 | 4 | 2012 | 149 | 1620.5 | - | Yes<sup>2</sup> | 2 |
| *Tringa nebularia* | Scotland | 58.533 | -4.232 | historic | 0.020 | 0.005 | 43 | 1993 | 71 | 918.3 | 0.5 | Yes<sup>1</sup> | 4, 5 |
|  |  |  |  | recent | 0.029 | 0.010 | 7 | 2004 | 24 | 275.9 | 0.5 | Yes | 5 |
| *Arenaria interpres* | Greenland | 74.478 | -20.555 | historic | 0.034 | 0.011 | 4 | 1998 | 38 | 338.5 | - | - | 6 |
|  |  |  |  | recent | 0.069 | 0.006 | 16 | 2008 | 150 | 1238.0 | - | - | 6 |
| *Philomachus pugnax* | Russia | 72.906 | 106.104 | historic | 0.075 | 0.009 | 6 | 1996 | 79 | 810.9 | 0.6 | - | 7 |
|  |  |  |  | recent | 0.061 | 0.005 | 12 | 2006 | 176 | 1952.3 | 0.6 | - | 2, 7 |
| *Calidris alba* | Greenland | 74.478 | -20.555 | historic | 0.015 | 0.007 | 4 | 1998 | 35 | 387.3 | - | Yes | 6 |
|  |  |  |  | recent | 0.052 | 0.008 | 7 | 2003 | 58 | 642.7 | - | Yes<sup>1</sup> | 6 |
| *Calidris mauri* | Alaska | 64.449 | -164.977 | historic | 0.020 | 0.002 | 6 | 1996 | 219 | 3184.5 | - | Yes<sup>1</sup> | 2 |
|  |  |  |  | recent | 0.033 | 0.003 | 6 | 2012 | 288 | 3767.0 | - | Yes<sup>1</sup> | 2 |
| *Calidris temminckii* | Finland | 65.021 | 24.72 | historic | 0.059 | 0.004 | 19 | 1992 | 464 | 3031.8 | 0.5 | Yes<sup>2</sup> | 8 |
|  |  |  |  | recent | 0.018 | 0.002 | 8 | 2004 | 153 | 2845.8 | 0.9 | Yes<sup>1</sup> | 9 |
| *Calidris melanotos* | Russia | 72.906 | 106.104 | historic | 0.075 | 0.004 | 6 | 1996 | 248 | 2675.4 | 0.6 | - | 7 |
|  |  |  |  | recent | 0.071 | 0.003 | 12 | 2006 | 364 | 4058.9 | 0.6 | - | 2, 7 |
| *Calidris melanotos* | Russia | 68.610 | 171.241 | historic | 0.032 | 0.011 | 9 | 1976 | 23 | 247.0 | 0.5 | - | 10 |
|  |  |  |  | recent | 0.055 | 0.024 | 3 | 2013 | 14 | 121.5 | - | - | 2 |
| *Calidris alpina* | Canada | 58.698 | -93.942 | historic | 0 | 0 | 4 | 1966 | 13 | 162.5 | 0.5 | Yes | 1 |
|  |  |  |  | recent | 0.023 | 0.003 | 4 | 2012 | 110 | 1493.5 | - | Yes<sup>3</sup> | 2 |
| *Calidris alpina* | Greenland | 74.478 | -20.555 | historic | 0.012 | 0.007 | 4 | 1998 | 28 | 332.1 | - | - | 6 |
|  |  |  |  | recent | 0.034 | 0.004 | 16 | 2008 | 184 | 2037.3 | - | - | 6 |
| *Calidris alpina* | Russia | 72.906 | 106.104 | historic | 0.059 | 0.006 | 6 | 1996 | 129 | 1335.0 | 0.6 | - | 7 |
|  |  |  |  | recent | 0.060 | 0.004 | 12 | 2006 | 180 | 2104.5 | 0.6 | - | 2, 7 |
| *Calidris alpina* | Russia | 68.610 | 171.241 | historic | 0.032 | 0.008 | 9 | 1976 | 51 | 506.2 | 0.5 | - | 10 |
|  |  |  |  | recent | 0.051 | 0.013 | 3 | 2013 | 45 | 388.0 | - | - | 2 |
| *Calidris minuta* | Russia | 72.906 | 106.104 | historic | 0.070 | 0.012 | 6 | 1996 | 49 | 477.8 | 0.6 | - | 7 |
|  |  |  |  | recent | 0.048 | 0.003 | 12 | 2006 | 228 | 2709.9 | 0.6 | - | 2, 7 |
| *Calidris minutilla* | Canada | 58.698 | -93.942 | historic | 0.010 | 0.004 | 4 | 1966 | 56 | 612.0 | 0.5 | - | 1 |
|  |  |  |  | recent | 0.003 | 0.004 | 3 | 2013 | 21 | 255.0 | - | - | 2 |
| *Calidris pusilla* | Alaska | 64.449 | -164.977 | historic | 0.031 | 0.004 | 6 | 1996 | 187 | 2273.5 | - | - | 2 |
|  |  |  |  | recent | 0.051 | 0.004 | 5 | 2012 | 213 | 2396.5 | - | - | 2 |
| *Calidris pusilla* | Alaska | 70.380 | -149.534 | historic | 0.012 | 0.002 | 4 | 1990 | 179 | 1962.2 | 0.5 | - | 11 |
|  |  |  |  | recent | 0.004 | 0.005 | 2 | 2012 | 21 | 303.0 | - | - | 2 |
| *Phalaropus lobatus* | Alaska | 64.449 | -164.977 | historic | 0.031 | 0.009 | 3 | 1994 | 46 | 476.0 | - | - | 2 |
|  |  |  |  | recent | 0.056 | 0.006 | 5 | 2012 | 149 | 1379.5 | - | - | 2 |
| *Phalaropus lobatus* | Russia | 68.610 | 171.241 | historic | 0.040 | 0.009 | 9 | 1976 | 52 | 455.6 | 0.5 | - | 10 |
|  |  |  |  | recent | 0.062 | 0.021 | 2 | 2013 | 16 | 133.0 | - | - | 2 |
| *Phalaropus fulicarius* | Russia | 72.906 | 106.104 | historic | 0.074 | 0.006 | 6 | 1996 | 135 | 1354.3 | 0.6 | - | 7 |
|  |  |  |  | recent | 0.044 | 0.003 | 12 | 2006 | 317 | 3395.6 | 0.6 | - | 2, 7 |

<sup>a</sup> Taxonomic order in the IOC World Bird List has changed recently, so we ordered species to follow Table S4 of Kubelka et al. for ease of comparison.
<sup>b</sup> B = value used in the Beintema transformation (see text) to calculate exposure days; shown only when the transformation was necessary.
<sup>c</sup> "Yes" indicates populations included in Kubelka et al. with the following caveats: 1) fewer years and nests, 2) fewer nests from the same years, 3) assumed all nests that failed to unknown causes were depredated. In some cases, Kubelka et al. also used a different value for B (see their supporting data for the corresponding value). Populations not included by Kubelka et al., all of which met their criteria for inclusion, are indicated with "-".
<sup>d</sup> Sources from Kubelka et al: 1) Jehl 1971, 2) Arctic Shorebird Demographics Network 2016, 3) Skeel 1983, 4) Christian & Hancock 2009, 5) Hancock in litt., 6) Hansen in litt., 7) Soloviev et al. 2010, 8) Rönkä et al. 2003, 9) Thompson et al. 2014, 10) Kondratyev 1982, 11) Moitoret et al. 1996.

**Table S8.** Daily predation rates in relation to period and Beintema conversion coefficient.

| B | β<sub>period</sub> | Intercept | β<sub>latitude</sub> |
| --- | --- | --- | --- |
| 0.5 | **0.50 (0.19)** | -5.17 (6.31) | 0.02 (0.10) |
| 0.4 | 0.31 (0.20) | -6.42 (6.28) | 0.04 (0.10) |
| 0.3 | 0.16 (0.22) | -6.67 (7.47) | 0.05 (0.10) |
| 0.2 | -0.07 (0.26) | -7.04 (6.69) | 0.06 (0.10) |
| 0.1 | -0.51 (0.33) | -7.72 (6.97) | 0.07 (0.11) |

Results of linear mixed-effects models testing for an effect of period on daily predation rates under various assumptions for Beintema coefficients (B = 0.5, 0.4, 0.3, 0.2, or 0.1). Values in parentheses are SEs; bold values indicate estimates significantly different from zero. Latitude was scaled by subtracting the mean and dividing by 1 SD.

## References

1. G.-R. Walther, E. Post, P. Convey, A. Menzel, C. Parmesan, T. J. C. Beebee, J.-M. Fromentin, O. Hoegh-Guldberg, F. Bairlein, Ecological responses to recent climate change. Nature 416, 389–395 (2002).

2. N. Jonzén, A. Lindén, T. Ergon, E. Knudsen, J. O. Vik, D. Rubolini, D. Piacentini, C. Brinch, F. Spina, L. Karlsson, M. Stervander, A. Andersson, J. Waldenström, A. Lehikoinen, E. Edvardsen, R. Solvang, N. Chr. Stenseth, Rapid advance of spring arrival dates in long-distance migratory birds. Science 312, 1959–61 (2006).

3. S. J. Thackeray, P. A. Henrys, D. Hemming, J. R. Bell, M. S. Botham, S. Burthe, P. Helaouet, D. G. Johns, I. D. Jones, D. I. Leech, E. B. Mackay, D. Massimino, S. Atkinson, P. J. Bacon, T. M. Brereton, L. Carvalho, T. H. Clutton-Brock, C. Duck, M. Edwards, J. M. Elliott, S. J. G. Hall, R. Harrington, J. W. Pearce-Higgins, T. T. Høye, L. E. B. Kruuk, J. M. Pemberton, T. H. Sparks, P. M. Thompson, I. White, I. J. Winfield, S. Wanless, Phenological sensitivity to climate across taxa and trophic levels. Nature 535, 241–245 (2016).

4. D. Zurell, C. H. Graham, L. Gallien, W. Thuiller, N. E. Zimmerman, Long-distance migratory birds threatened by multiple independent risks from global change. Nature Climate Change 8, 992–996 (2018).

5. V. Kubelka, M. Šálek, P. Tomkovich, Z. Végvári, R. P. Freckleton, T. Székely, Global pattern of nest predation is disrupted by climate change in shorebirds. Science 362, 680–683 (2018).

6. C. E. Studds, B. E. Kendall, N. J. Murray, H. B. Wilson, D. I. Rogers, R. S. Clemens, K. Gosbell, C. J. Hassell, R. Jessop, D. S. Melville, D. A. Milton, C. D. T. Minton, H. P. Possingham, A. C. Riegen, P. Straw, E. J. Woehler, R. A. Fuller, Rapid population decline in migratory shorebirds relying on Yellow Sea tidal mudflats as stopover sites. Nat. Commun. 8, 1–7 (2017).

7. M. Bulla, J. Reneerkens, E. L. Weiser, R. B. Lanctot, B. Kempenaers, Supporting information for comment on “Global pattern of nest predation is disrupted by climate change in shorebirds.” Open Science Framework, https://osf.io/x8fs6/

8. H. Mayfield, Nesting success calculated from exposure. Wilson bull. 73, 255–261 (1961).

9. H. Mayfield, Suggestions for calculating nest success. Wilson bull. 87, 456–466 (1975).

10. A. J. Beintema, Inferring nest success from old records. Ibis 138, 568–570 (1996).

11. S. J. Dinsmore, G.C. White, F.L. Knopf, Advanced techniques for modeling avian nest survival. Ecology 83, 3476–3488 (2002).

12. T. L. Shafer, F.R. Thompson, III, Making meaningful estimates of nest survival with model-based methods. Studies in Avian Biology 34, 84–95 (2007).

13. J. J. Rotella, Modeling nest-survival data: recent improvements and future directions. Studies in Avian Biology 34, 145–148 (2007).

14. N. Verboven, B. J. Ens, S. Dechesne, Effect of investigator disturbance on nest attendance and egg predation in Eurasian oystercatchers. Auk 118, 503–508.

15. B. W. Meixell, P. L. Flint, Effects of industrial and investigator disturbance on Arctic-nesting geese. J. Wildl. Manag. 81, 1372–1385 (2017).

## References

1 R Core Team. R: A language and environment for statistical computing - version 3.5.1. Internet. R Foundation for Statistical Computing [cited 2019 Jan 25], http://www.R-project.org/ (2018).

2 Therneau, T. M. coxme: Mixed Effects Cox Models. R package version 2.2-10. https://CRAN.R-project.org/package=coxme (2018).

3 Kubelka, V. et al. Global pattern of nest predation is disrupted by climate change in shorebirds. Science 362, 680–683 (2018).

4 Bates, D., Maechler, M., Bolker, B. & Walker, S. Fitting Linear Mixed-Effects Models Using lme4. J Stat Softw 67, 1–48 (2015).

5 Gelman, A. & Hill, J. Data analysis using regression and multilevel/hierarchical models (Cambridge University Press, 2007).

6 Gelman, A. & Su, Y.-S. arm: Data Analysis Using Regression and Multilevel/Hierarchical Models. R package version 1.8-6. http://CRAN.R-project.ore/packaee=arm (2015).

7 R-Core-Team. R: A Language and Environment for Statistical Computing. Version 3.3.0. R Foundation for Statistical Computing, http://www.R-project.org/ (2016).

8 Bates, D., Maechler, M., Bolker, B. & Walker, S. lme4: Linear mixed-effects models using Eigen and S4. R Package Version 1.0-6. http://CRAN.R-project.ore/packaee=lme4 (2014).

9 Mayfield, H. Nesting success calculated from exposure. Wilson Bull 73, 255–261 (1961).

10 Beintema, A. J. Inferring nest success from old records. Ibis 138, 568–570 (1996).

11 Lanctot, R. B., S. C. Brown, and B. K. Sandercock. Data from: Arctic Shorebird Demographics Network. NSF Arctic Data Center (2016). https://arcticdata.io/cataloe/view/doi:10.18739/A2CD5M

12 Brown, S. C., H. R. Gates, J. R. Liebezeit, P. A. Smith, B. L. Hill, and R. B. Lanctot. Arctic Shorebird Demographics Network Breeding Camp Protocol, Version 5. U.S. Fish and Wildlife Service and Manomet Center for Conservation Sciences (2014).

13 Johnson, D. H. Estimating nest success: The Mayfield method and an alternative. Auk 96, 651–661 (1979).

